# Mapping the landscape of genetic dependencies in chordoma

**DOI:** 10.1101/2022.08.17.504193

**Authors:** Tanaz Sharifnia, Mathias J. Wawer, Amy Goodale, Yenarae Lee, Mariya Kazachkova, Joshua M. Dempster, Sandrine Muller, Joan Levy, Daniel M. Freed, Josh Sommer, Jérémie Kalfon, Francisca Vazquez, William C. Hahn, David E. Root, Paul A. Clemons, Stuart L. Schreiber

## Abstract

Identifying the spectrum of genes required for cancer cell survival can reveal essential cancer circuitry and therapeutic targets, but such a map remains incomplete for many cancer types. We applied genome-scale CRISPR-Cas9 loss-of-function screens to map the landscape of selectively essential genes in chordoma, a bone cancer with few validated targets. This approach confirmed a known chordoma dependency, *TBXT* (*T*; brachyury), and identified a range of additional dependencies, including *PTPN11, ADAR, PRKRA, LUC7L2, SRRM2, SLC2A1, SLC7A5, FANCM*, and *THAP1. CDK6, SOX9, and EGFR*, genes previously implicated in chordoma biology, were also recovered. We found genomic and transcriptomic features that predict specific dependencies, including interferon-stimulated gene expression, which correlates with *ADAR* dependence and is elevated in chordoma. Validating the therapeutic relevance of dependencies, small-molecule inhibitors of SHP2, encoded by *PTPN11*, had potent preclinical efficacy against chordoma. Our results generate an emerging map of chordoma dependencies to enable biological and therapeutic hypotheses.

## MAIN

Elucidating the set of genes upon which cancer cell proliferation and survival is especially reliant (the ‘dependencies’ of cancer) can provide an important roadmap for the treatment and characterization of cancer. Knowledge of these vulnerabilities can guide the choice of cancer-selective drugs, reveal mechanisms of cancer cell initiation or maintenance, and permit the classification of cancer subtypes or cell states^1^. The observation that clinically relevant dependencies can be predicted by somatic mutation or copy-number alteration of the same gene (exemplifying the phenomenon of ‘oncogene addiction’) has motivated large-scale efforts to sequence tumor genomes and thereby facilitate cancer-dependency discovery^2,3^. However, identifying tractable therapeutic vulnerabilities using tumor genome sequencing has been more challenging for cancer types that harbor relatively few somatic mutations, when gene mutations that enhance fitness cannot be readily distinguished from bystander mutations, or when altered genes cannot be modulated with the existing arsenal of drugs.

Alongside efforts to identify somatically mutated cancer-dependency genes, a growing number of tumor-intrinsic cancer dependencies have been discovered via functional approaches whose classification extends beyond that of canonical mutated oncogenes^1,4^. These include dependencies reflecting synthetic lethal interactions^5,6^, such as those occurring when loss of one member of a paralog pair leads to increased reliance on the other member (‘paralog dependencies’)^7,8^, or those resulting from hemizygous copy-number loss and/or reduced expression of the same gene (‘CYCLOPS’ genes)^9^; dependencies arising from dysregulated or persistent expression of master regulatory genes that mediate normal lineage development (‘lineage-survival’ dependencies)^10^; and dependencies unique to a particular cancer cell state^11^. In this way, many cancer dependencies can be classified as ‘non-oncogene’ dependencies, whereby the unique molecular features of the cancer confer a heightened dependence on the normal cellular function of specific genes^12^.

Indeed, dependencies resulting from somatically mutated oncogenes now appear to represent only a minority of cancer dependencies overall^4^. Notably, cancer types that harbor relatively few somatic mutations, such as some pediatric cancers, do not necessarily have a reduced number of genetic dependencies relative to cancer types with a higher mutational burden^13^. Together, these observations suggest the existence of a spectrum of cancer dependencies that may not be discoverable via tumor genome sequencing, but which may nevertheless serve as effective therapeutic targets. Yet for many cancer types, a comprehensive map of differential dependencies from which such targets could be nominated remains incomplete.

One such cancer type is chordoma, a primary bone cancer for which there is a limited number of validated cancer targets and no approved systemic therapy^14^. Chordoma is a genomically quiet cancer: genomic sequencing of chordoma tumors has demonstrated that nearly half of all sporadic chordoma cases do not harbor any known driver mutation, and most somatic alterations that do exist are not currently targetable^15,16^ -- limiting the ability to identify an appropriate targeted therapy in most patients using traditional genome-guided precision medicine approaches. Nonetheless, chordoma tumor cells may have distinct genomic or functional features that render them especially dependent on the activities of specific genes. We sought to identify vulnerabilities that are conferred by the unique cellular circuitry of chordoma, with a view to revealing novel approaches for the treatment of this disease.

To this end, here we applied genome-scale CRISPR-Cas9 loss-of-function screens to map the landscape of selective dependencies in chordoma. Previously, CRISPR-Cas9 screens in two chordoma cell lines had identified the developmental transcription factor *T* (brachyury), renamed *TBXT*, to be the top selectively essential gene in chordoma, relative to 125 non-chordoma cancer cell lines^17^. While *TBXT* had been the only statistically significant dependency gene identified, we hypothesized that this could be attributed to the small set of chordoma cell lines screened, and that the full range of tumor dependencies in chordoma remained to be discovered. Moreover, *TBXT* encodes a transcription factor that is currently considered challenging to drug, further motivating a search for additional dependency genes.

In this study, by expanding the set of cell lines subjected to genome-scale loss-of-function screening, we recovered *TBXT* and further discovered a spectrum of previously unappreciated chordoma dependency genes. We demonstrated that targeting one such dependency in preclinical models of chordoma resulted in profound antitumor efficacy, providing a rationale for new clinical trials in chordoma. Together, these findings generate an emerging map of selectively essential genes in chordoma, facilitating future studies that seek to understand the unique tumor biology of this cancer type.

## RESULTS

### Genome-scale CRISPR-Cas9 screens identify a spectrum of selectively essential genes in chordoma

To map the landscape of genes essential for chordoma cell viability, we performed genome-scale pooled CRISPR-Cas9 loss-of-function screens in two chordoma cell lines (JHC7, U-CH2), thereby doubling the number of chordoma cell lines profiled using this approach^17^. A library of >74,000 single-guide RNAs (sgRNAs) targeting ∼18,560 genes (Methods) was introduced via lentiviral transduction into stably Cas9-expressing chordoma cells. Cells were grown in culture for 20-22 d, after which sgRNAs were quantified from the genomic DNA (gDNA) of surviving cells using massively parallel sequencing. SgRNAs that were depleted from the cell population relative to the screening library were inferred to target candidate essential genes. Gene-dependency scores were generated by each of two established approaches: directly quantifying the degree to which the loss of a gene impacts cell viability (CERES score), and further transforming this value into a probability of gene dependency (probability score)^18,19^. To identify dependencies selective for chordoma, and remove commonly essential genes, we compared gene-dependency scores derived from four chordoma cell lines (JHC7, U-CH2, UM-Chor1, and MUG-Chor1) to those from 765 non-chordoma cancer cell lines similarly subjected to CRISPR-Cas9 loss-of-function screens as part of the Broad Institute Cancer Dependency Map project (https://depmap.org/portal/)^18^ (**Figure 1a**).

**Figure 1.**
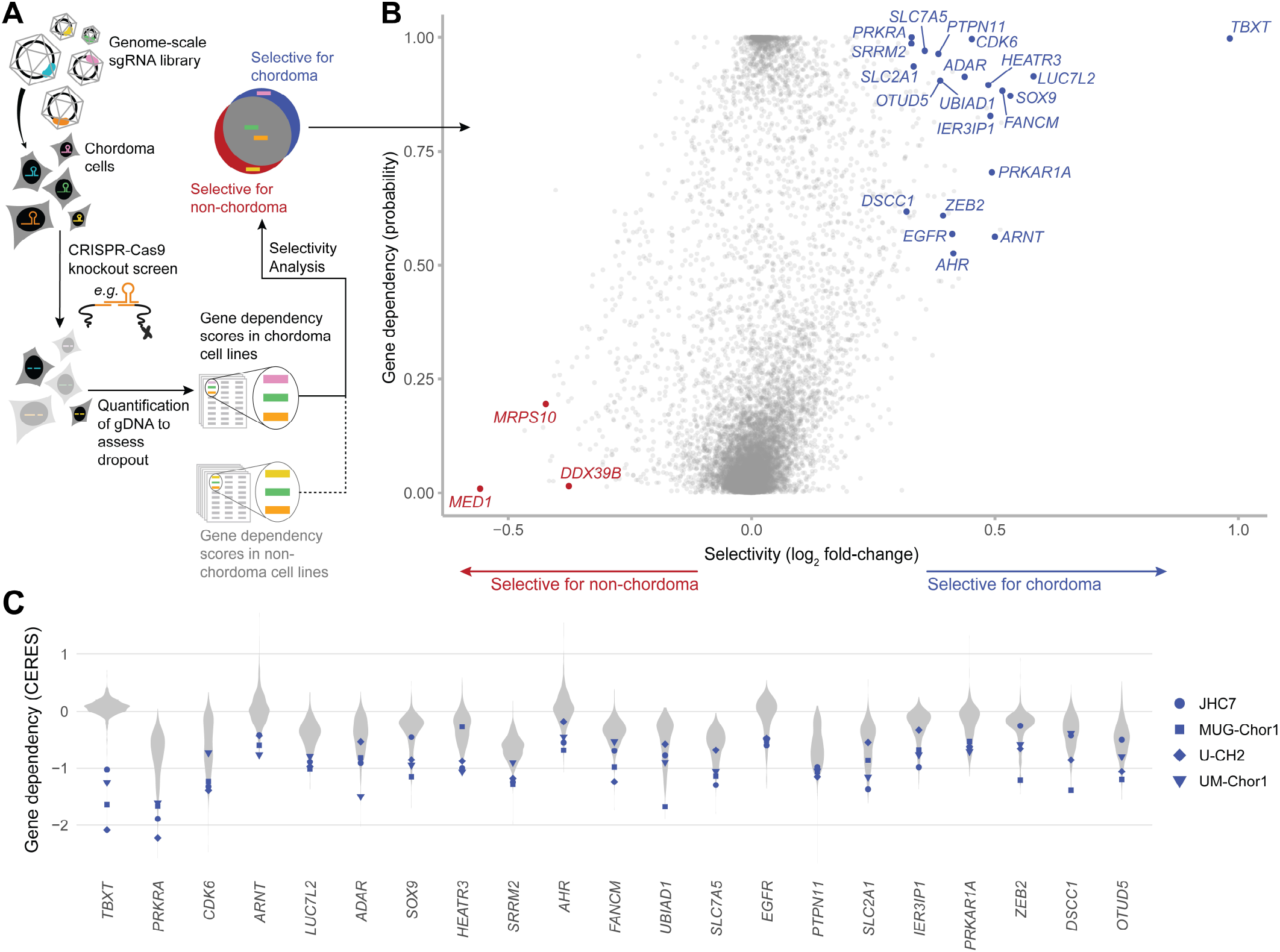
Genome-scale CRISPR-Cas9 screens identify a spectrum of selectively essential genes in chordoma. **A)** Experimental workflow of genome-scale, loss-of-function CRISPR-Cas9 screens to identify selectively essential genes in chordoma. **B)** Selective essentiality analysis identifying chordoma dependency genes. Selectivity is quantified by the log2 fold-change in mean gene dependency (using probability scores) between four chordoma and 765 non-chordoma cell lines. Gene dependencies selective for chordoma/non-chordoma are indicated in blue/red (see Methods for details). **C)** Distribution of gene dependency (using CERES scores) for the indicated genes across four chordoma cell lines (blue) and 765 non-chordoma cell lines (gray; non-chordoma cell lines were profiled as part of the Broad Institute Cancer Dependency Map). Genes are ranked by decreasing selectivity for chordoma, quantified by the log2 fold-change in mean CERES scores between four chordoma and 765 non-chordoma cell lines.

Consistent with previous findings, the top selectively essential gene, out of over 18,000 analyzed, was *TBXT* (**Figure 1b, Supplementary Figure 1**)^17^. In addition to *TBXT*, this analysis identified a diverse spectrum of selectively essential genes with previously unappreciated relevance to chordoma, including *PTPN11, ADAR, PRKRA, LUC7L2, SRRM2, SLC2A1, SLC7A5, FANCM, AHR, ARNT, HEATR3, UBIAD1, IER3IP1, PRKAR1A, ZEB2, DSCC1*, and *OTUD5* (**Figure 1b-c, Supplementary Figure 1**). Genes previously implicated in chordoma biology, including *CDK6, SOX9*, and *EGFR*, were also recovered^20-22^. Conversely, a small number of genes was identified whose gene-dependency scores were of a greater magnitude in non-chordoma than in chordoma cell lines, and these genes included *MED1, MRPS10*, and *DDX39B* (**Figure 1b, Supplementary Figure 1**).

To classify and prioritize candidate essential genes in chordoma, we first asked whether these genes could be grouped into shared biological pathways or functions. Functionally related genes often exhibit similar patterns of essentiality across cell lines^23-25^; thus, we measured whether candidate dependency genes display correlated patterns of essentiality across 769 cancer cell lines that had been subjected to CRISPR-Cas9 loss-of-function screening (Broad Institute Cancer Dependency Map, https://depmap.org/portal/; **Figure 2a**). Using this approach, four groups of functionally related genes emerged: (1) *FANCM, SRRM2, ZEB2*, and *DSCC1*; (2) *UBIAD1, PRKRA*, and *ADAR*; (3) *AHR, ARNT*, and *TBXT*; and (4) *PTPN11* and *EGFR* (**Figure 2a**). Most similar were gene pairs encoding proteins with known functional relationships: *AHR* and *ARNT* form a transcriptionally active heterodimer^26^; *FANCM* and *DSCC1* each has roles in various DNA replication-related processes^27,28^; *PTPN11* can act as a positive effector of mitogenic signaling induced by the receptor tyrosine kinase *EGFR*^29^; and *ADAR* and *PRKRA* regulate interferon responses^30^. Complementing the co-essentiality analysis, we also grouped dependency genes based on publicly available protein-protein interaction annotations (**Figure 2b**). This analysis yielded a single cluster of genes comprising *EGFR, PRKAR1A, PTPN11, SOX9, ZEB2, SLC7A5, SLC2A1, ARNT*, and *AHR* (**Figure 2b**) and recovered known strong connections for *PTPN11*/*EGFR* and *AHR*/*ARNT*. Several candidate genes, including *OTUD5, CDK6, LUC7L2, IER3IP1*, and *HEATR3* were not linked to other chordoma dependency genes following either of these two approaches (**Figure 2a-b**).

**Figure 2.**
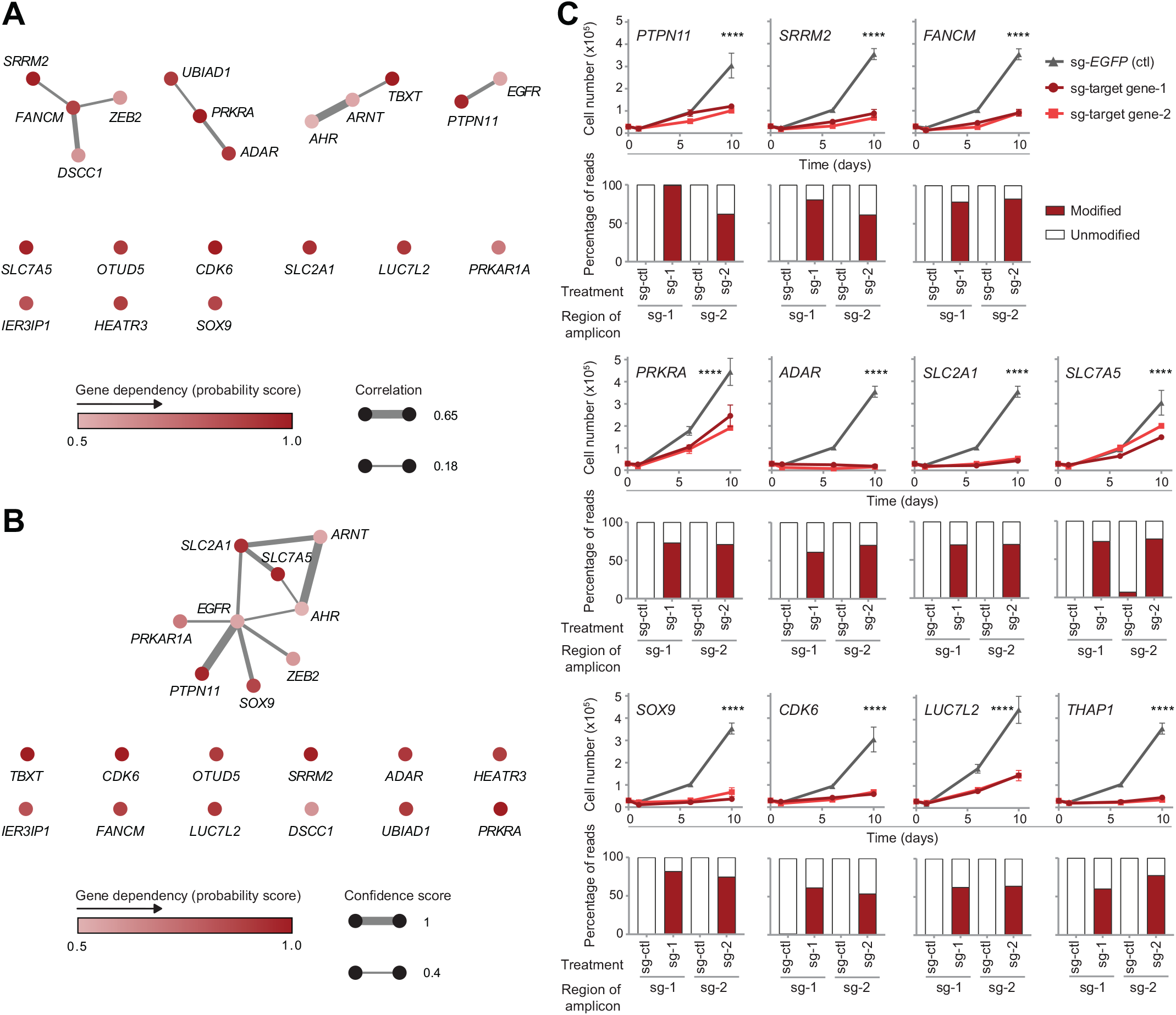
Validation of candidate chordoma dependency genes. **A)** Co-essentiality network for selective chordoma dependency genes. Nodes: chordoma dependency genes, colored by dependency probability scores. Edges: Pearson correlation ≥ 0.18 between gene-dependency profiles, *i*.*e*., dependency probability scores across all 769 cancer cell lines; edge width scaled by correlation coefficient. B) STRING protein-protein interaction network for selective chordoma dependency genes. Nodes: chordoma dependency genes, colored by dependency probability scores. Edges: putative interactions with a STRING confidence score ≥ 0.4; edge width scaled by confidence score. **C)** (Top rows) Proliferation of Cas9-expressing UM-Chor1 chordoma cell lines transduced with one of two distinct sgRNAs targeting a candidate dependency gene or a non-targeting sgRNA control. Points represent the mean ± s.d. (*n* = 3 biological samples measured in parallel). \*\*\*\**P* < 0.0001, derived from a two-way analysis of variance (ANOVA). *P* values for the test comparing sg-*EGFP* and sg-target gene*-*1 are displayed and refer to the time × treatment interaction. Graphs with identical sg-*EGFP* control curves reflect experiments performed in parallel on the same day. (Bottom rows) Percentage of modified versus unmodified reads following amplicon sequencing of the indicated sgRNA-containing genomic site following sgRNA treatment of UM-Chor1-Cas9 cells (sg-*EGFP* non-targeting control or one of two distinct sgRNAs targeting a candidate dependency gene). The small fraction of “modified” reads observed for the sg-*SLC7A5*-2 amplicon following sg-*EGFP* control treatment originates from a residual amount of single-nucleotide variation we were unable to match to genomically defined off-target amplicons. However, their distribution points to an additional off-target amplicon as the source (rather than true editing events).

We applied this pathway analysis to select a diverse subset of candidate dependency genes for validation and further functional characterization. Genes were selected to represent both those functionally connected to other chordoma dependency genes (*PTPN11, FANCM, ADAR, PRKRA, SOX9, SRRM2, SLC2A1, SLC7A5*), as well as those with putatively unique effects (*LUC7L2, CDK6*). We included an additional gene for validation, *THAP1*, that is essential across all cancer cell lines, yet whose loss has a greater degree of viability reduction in chordoma cell lines compared to non-chordoma cell lines (**Supplementary Figure 1**).

To validate the essentiality of a candidate dependency gene, Cas9-expressing UM-Chor1 chordoma cells were transduced in parallel with two independent sgRNAs targeting a candidate dependency gene and a non-targeting control and assayed for cell viability (**Figure 2c**). Consistent with primary screening results, sgRNA-mediated gene suppression of all candidate dependency genes tested led to impaired chordoma cell proliferation (**Figure 2c**). SgRNA-mediated gene editing was confirmed for all validated genes, via amplicon sequencing of treated cells (**Figure 2c**); and sgRNA-mediated protein suppression was measured and confirmed for a subset of validated genes (**Supplementary Figure 2**). The high validation rate of the candidate dependency genes tested demonstrates reliability of the primary screening analysis to identify *bona fide* chordoma dependency genes, which likely extend beyond those chosen for functional follow-up.

### Genomic and transcriptomic features that predict specific gene dependencies

Identifying a molecular feature that is associated with a specific genetic dependency can help classify tumors by their expected response to a targeted therapy and reveal the mechanisms underlying gene essentiality. To explore genomic and transcriptomic features that predict specific gene dependencies discovered in our screens, we queried gene-mutation, gene-copy-number, and gene-expression data generated from over 700 cancer cell lines as part of the Cancer Cell Line Encyclopedia (CCLE)^31^. For each chordoma dependency gene, we correlated its signature of essentiality across all cancer cell lines in the Broad Institute Cancer Dependency Map with these genomic and transcriptomic profiling data.

This analysis recovered mutations in genes known to act in the same pathway as specific dependency genes: decreased dependence on *CDK6* can be predicted by nonsense and splice site mutations in the downstream target *RB1*^32^; decreased dependence on *PTPN11* can be predicted by missense mutations in the downstream effectors *BRAF, NRAS*, or *KRAS*^4,33^; and increased dependence on *FANCM* can be predicted by splice site mutations in its functional and synthetic lethal partner, *BRCA1*^34^. (**Figure 3a**).

**Figure 3.**
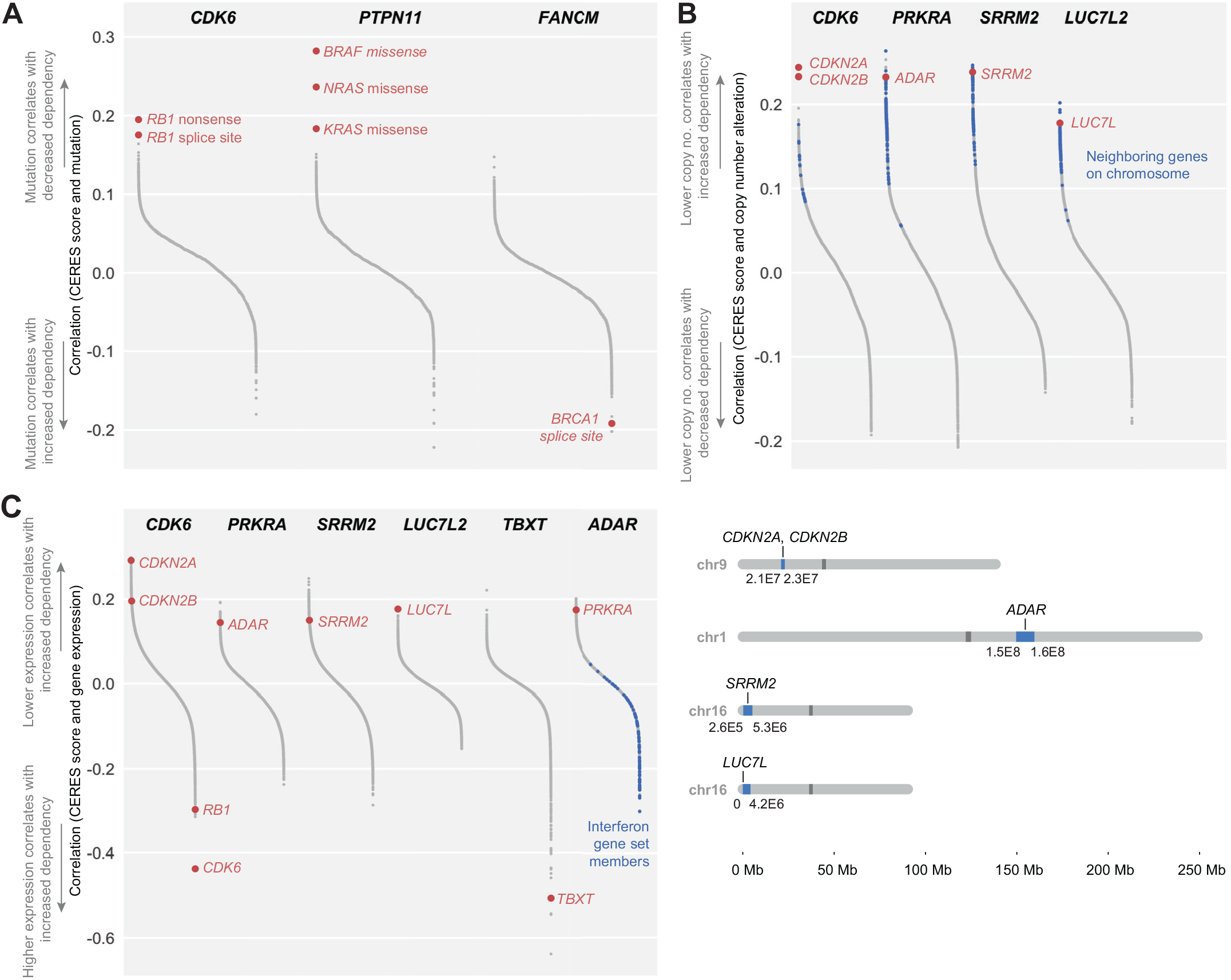
Genomic and transcriptomic predictors of gene dependencies. **A)** Mutation correlates of selected dependency genes. **B)** (Top) Copy-number correlates of selected dependency genes. Blue: genes neighboring the labeled correlate on the same chromosome, within a window selected for maximum enrichment of neighboring genes at the top of the correlation rank list (see Methods). (Bottom) Genomic loci of copy-number correlates and window selected by enrichment analysis. **C)** Gene-expression correlates of selected dependency genes. Members of the MSigDB Interferon Alpha gene set are indicated in blue. For all calculations, we used pairwise-complete observations to handle occasional missing values. All comparisons included at least 719 cell lines, and over 95% of comparisons included the complete set of cell lines for which data were available (mutations: 768; gene expression: 767; copy number: 769).

We also found copy-number changes in several categories of genes to be associated with specific gene dependencies (**Figure 3b**). These included copy-number changes in genes known to act in the same pathway as the dependency gene, such as the previously reported correlation between lower copy number of *CDKN2A*/*CDKN2B* with CDK4/6 dependency^35^; as well as the correlation between *ADAR* copy-number alteration with *PRKRA* dependency. In the case of *SRRM2*, copy-number changes correlate with dependency on the same gene (*SRRM2*). Lastly, this analysis identified copy-number alteration in the genetic paralog of a dependency gene: lower copy number of *LUC7L* is predictive of *LUC7L2* dependency (**Figure 3b**).

As expected, correlations between gene dependency and copy-number alterations were similarly reflected at the gene-expression level: dependency on *CDK6, PRKRA, SRRM2*, and *LUC7L2* each could be predicted by gene-expression changes in *CDKN2A*/*CDKN2B*; *ADAR*; *SRRM2*; and *LUC7L*, respectively (**Figure 3c**). *CDK6* dependency could also be predicted by higher gene expression of *RB1* and *CDK6* (**Figure 3c**). We also observed high *TBXT* expression to predict dependence on *TBXT*. This analysis also identified expression of interferon (IFN) genes to be correlated with dependency on the RNA adenosine deaminase *ADAR*, consistent with the observation that high expression levels of interferon-stimulated genes (ISGs) are a biomarker of *ADAR* dependency^36,37^.

Several of the correlated features identified using the set of over 700 cancer cell lines are present in individual chordoma cell lines and may have therapeutic relevance. For example, U-CH2 and MUG-Chor1 chordoma cell lines each show biallelic loss of *CDKN2A*, as well as high selective dependence on *CDK6* (**Figure 1c**). As somatic homozygous deletion of *CDKN2A* is a recurrent feature of chordoma tumors^15,16,38^, these results suggest that patients whose tumors harbor *CDKN2A* loss may benefit from inhibitors targeting CDK6. In addition to generating therapeutic hypotheses, we also asked whether correlated features could provide insight into the cellular circuitry of chordoma. As ISG expression levels have not previously been assessed in chordoma, we further investigated this feature in chordoma models.

### Interferon-stimulated genes are overexpressed in chordoma cells and further upregulated following *ADAR* gene suppression

Given the association of elevated ISG expression with *ADAR* dependence, and chordoma cells’ selective dependence on *ADAR* relative to non-chordoma cancer cell lines, we compared the expression of ISGs in chordoma cells versus that of other cancer types using a previously described 38-gene signature quantifying IFN pathway engagement (‘ISG core score’)^37^. Strikingly, chordoma cell lines have a higher median ISG core score than any of the 29 cancer lineages represented by at least two cell lines in the CCLE (**Figure 4a**), suggesting exceptionally high IFN signaling in chordoma.

**Figure 4.**
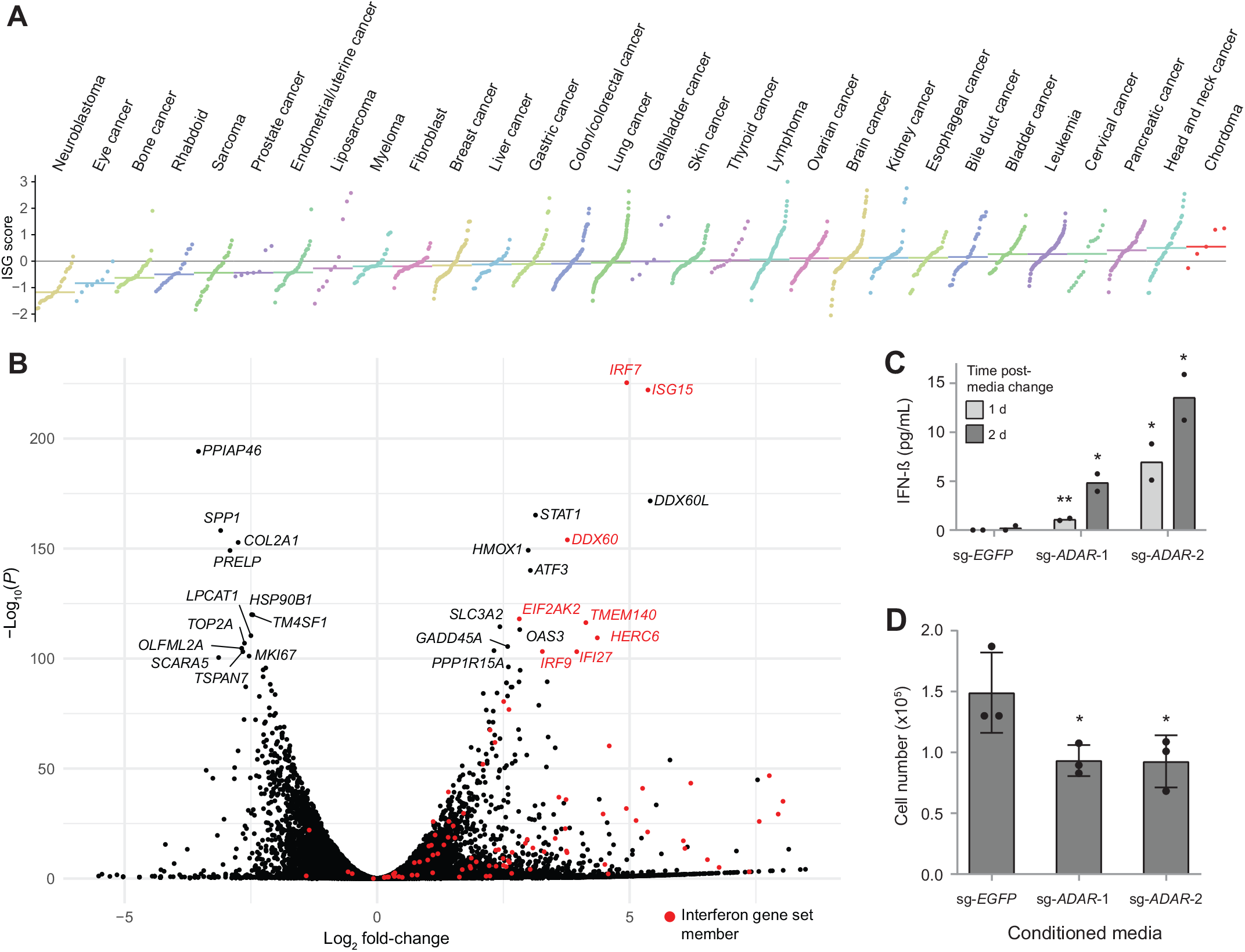
Interferon-stimulated genes are overexpressed in chordoma cells and further upregulated following *ADAR* gene suppression. **A)** Distribution of ISG core scores^37^ for chordoma cell lines and 1,294 non-chordoma cancer cell lines in the CCLE, grouped by lineage annotation. Colored horizontal bars: median scores for each group. Gray horizontal line: zero-score mark. **B)** Differential gene expression comparing the average effects of two distinct *ADAR*-targeting sgRNAs to a non-targeting sgRNA control in Cas9-expressing UM-Chor1 cells. Gene expression was measured with RNA sequencing. Members of the MSigDB Interferon Alpha gene set are indicated in red. **C)** IFN-β levels in conditioned media harvested from Cas9-expressing UM-Chor1 cells transduced with the indicated sgRNAs and subsequently subjected to a media change after selection for infected cells. IFN-β levels were measured by ELISA. Data represent the mean of two technical replicates. \**P* < 0.05, \*\**P* < 0.01, derived from a one-tailed, unpaired *t* test. The statistical test was performed on the indicated condition and the corresponding sg-*EGFP* control. D) Cell viability of parental UM-Chor1 cells treated for 3 d with conditioned media harvested from Cas9-expressing UM-Chor1 cells transduced with the indicated sgRNAs. Data represent the mean ± s.d. (*n* = 3 biological samples measured in parallel). \**P* < 0.05, derived from a one-tailed, unpaired *t* test. The statistical test was performed on the indicated condition and the sg-*EGFP* control.

In other cancer types, *ADAR* suppression in *ADAR*-dependent cells triggers production of type I IFNs and activation of the nucleic acid sensor PKR, a mediator of IFN-dependent growth arrest^36,37,39^. Consistent with these findings, interferon alpha and interferon gamma response genes are the most enriched among upregulated genes following sgRNA-mediated repression of *ADAR* in chordoma cells using both gene-set enrichment analysis (GSEA)^40^ (**Supplementary Figure 3a**), as well as a network-based enrichment method (GeLiNEA)^41^ (**Supplementary Figure 3b-c**). We observed upregulation of specific IFNs and ISGs, such as *IRF7* and *ISG15*; as well as *EIF2AK2*, which encodes PKR (**Figure 4b**).

IRF7 is a key regulator of type I IFN gene expression: it activates IFN-β expression and secretion, which in turn stimulates type I IFN receptor to further amplify *IRF7* expression via a positive feedback loop^42,43^. We assayed IFN-β secretion induced by *ADAR* depletion by measuring IFN-β levels in the conditioned media of sg-*ADAR*-treated chordoma cells. Consistent with the observed upregulation of *IRF7* expression, *ADAR* gene suppression induced secretion of IFN-β (**Figure 4c**). Furthermore, the conditioned media from *ADAR*-deficient cells was sufficient to reduce proliferation of parental chordoma cells, suggesting that paracrine mechanisms contribute to sg-*ADAR*-induced lethality and can act independently of *ADAR* gene suppression (**Figure 4d**). Taken together, these results indicate that chronic ISG expression is exceptionally pronounced in chordoma cells relative to other cancer types, and that *ADAR* gene suppression further upregulates these genes and induces the secretion of factors, such as IFN-β, which are sufficient to reduce chordoma cell viability in a non-cell autonomous fashion.

### Inhibitors of SHP2, encoded by *PTPN11*, represent new candidate therapeutic agents in chordoma

In addition to providing insights into the cellular circuitry of chordoma, identifying a range of chordoma dependency genes may reveal new therapeutic targets. Several chordoma dependency genes identified in the primary screens, including *PTPN11, EGFR*, and *CDK6*, encode proteins that are currently targetable with small-molecule inhibitors. We investigated the immediate therapeutic relevance of newly identified chordoma dependency genes by determining whether small-molecule inhibitors targeting these proteins have antiproliferative activity in chordoma cells. EGFR-inhibitor and CDK4/6-inhibitor sensitivity in chordoma has been described previously^20,21^; thus, we focused on inhibitors of Src homology-2 domain-containing phosphatase 2 (SHP2), the protein encoded by *PTPN11*. SHP2 is a tyrosine phosphatase that promotes signal transduction downstream of receptor tyrosine kinases to activate the RAS/mitogen-activated protein kinase (MAPK) cascade, and has additional functions in modulating tumor immunity^33^; the recent development of selective, allosteric compounds targeting SHP2 led us to investigate their therapeutic potential in chordoma^44-47^.

Six chordoma cell lines each were treated with allosteric compounds targeting SHP2 (RMC-4550 or SHP099) and assayed for cell viability (**Figure 5a-b**). For comparison, we also tested the *BRAF*^V600E^-mutant A2058 melanoma cell line and the MDA-MB-468 breast adenocarcinoma cell line, which are reported to be insensitive and sensitive to SHP2 inhibition, respectively^46^, and which accordingly exhibit differential dependence on *PTPN11* gene suppression (**Supplementary Figure 4**). All cell lines were subjected to both 14-d colony formation and 6-d multipoint concentration-response assays, the former of which better distinguished the sensitivity of control cell lines (**Figure 5a-b**). We observed that five of the six chordoma cell lines tested were sensitive to SHP2 inhibition, to a degree exceeding that observed for the MDA-MB-468 reference cell line (**Figure 5a-b**). We confirmed on-target activity for SHP2 inhibitors by detecting reduced phosphorylation of ERK 1/2 in sensitive cell lines, with RMC-4550 exhibiting more potent on-target activity than SHP099 (**Figure 5c**).

**Figure 5.**
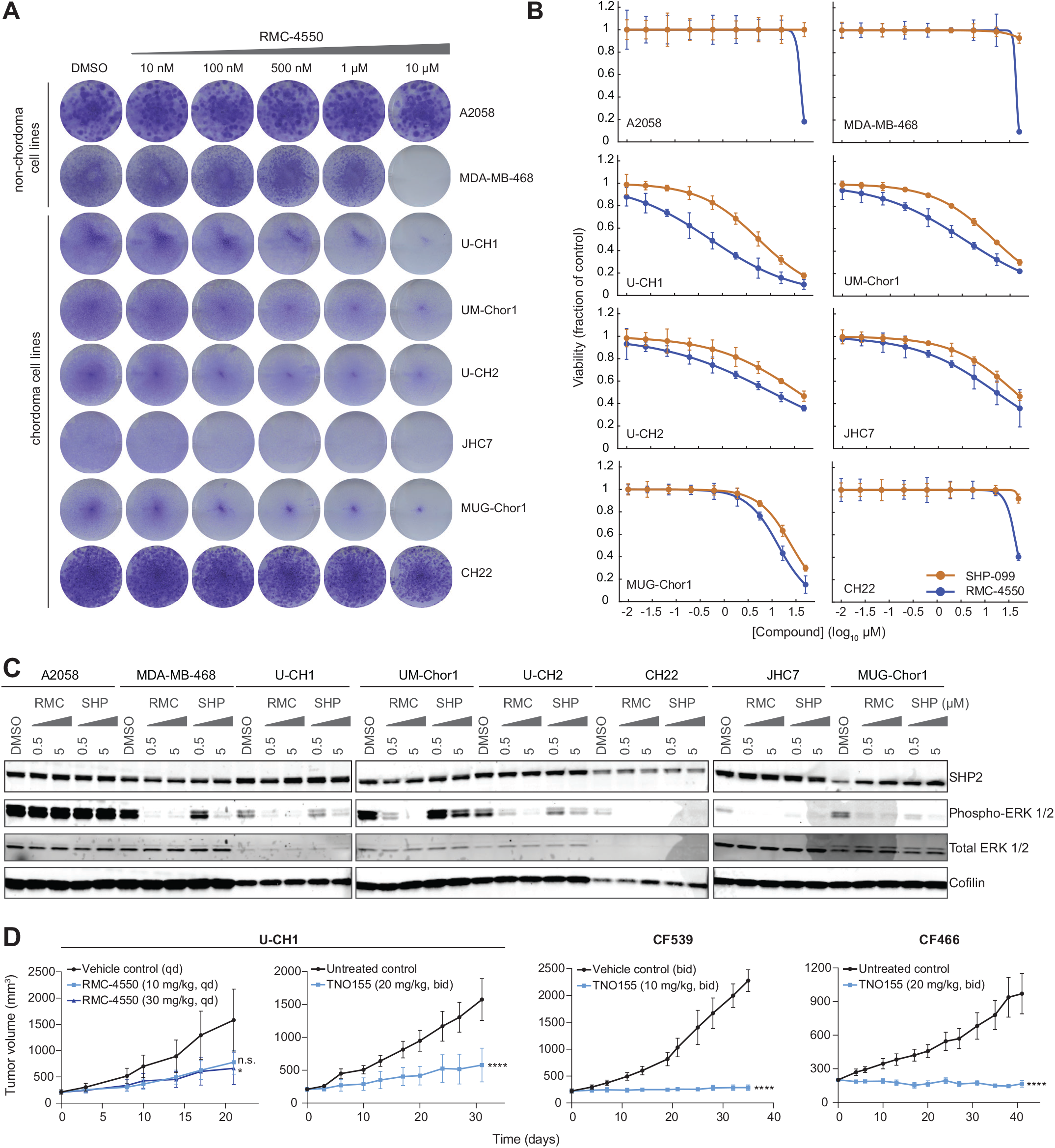
Inhibitors of SHP2, encoded by *PTPN11*, represent new candidate therapeutic agents against chordoma. **A)** Colony formation assays of chordoma and non-chordoma (negative control A2058; positive control MDA-MB-468) cell lines treated with indicated concentrations of RMC-4550 for 14 d. **B)** Viability of chordoma and non-chordoma (negative control A2058; positive control MDA-MB-468) cell lines treated with indicated concentrations of SHP2 inhibitors RMC-4550 and SHP099 and assayed for cell viability after 6 d with CellTiter-Glo. Response data are represented by a fitted curve to the mean fractional viability at each concentration relative to vehicle-treated cells; error bars represent the s.e.m. (*n* = 4 biological samples measured in parallel). **C)** Immunoblot analysis of chordoma and non-chordoma (negative control A2058; positive control MDA-MB-468) cell lines treated with indicated concentrations of RMC-4550, SHP099, or DMSO for 2 h. **D)** Tumor proliferation in mice engrafted with chordoma cells (U-CH1 cell line-derived xenograft, CF539 PDX, or CF466 PDX) and treated with a SHP2 inhibitor (RMC-4550 or TNO155). Points represent the mean tumor volume ± s.e.m. (*n* = 4-5 tumors for each arm of the U-CH1/RMC-4550 study; 6-7 tumors for each arm of the U-CH1/TNO155 study; 6-7 tumors for each arm of the CF539 study; 7 tumors for each arm of the CF466 study). n.s., not significant, \**P* < 0.05, \*\*\*\**P* < 0.0001, derived from a two-way analysis of variance (ANOVA) with repeated measures. *P* values for the time × treatment interaction (relative to the control condition) are indicated.

To test the efficacy of SHP2 inhibitors *in vivo*, a U-CH1-derived xenograft mouse model of chordoma was treated with 10 mg/kg or 30 mg/kg RMC-4550 or vehicle. The selected doses of RMC-4550 were determined to be capable of reducing phosphorylation of ERK 1/2 in the tumors of treated animals (**Supplementary Figure 5**). Consistent with *ex vivo* findings, when dosed for 21 days RMC-4550 inhibited tumor growth compared to vehicle treatment (**Figure 5d, Supplementary Figure 6**), with the 30 mg/kg dose meeting a threshold of significance (*P* = 0.044 for 30 mg/kg; *P* = 0.060 for 10 mg/kg). Alongside experiments with the tool compound RMC-4550, we tested a clinical-stage SHP2 inhibitor in this model to further evaluate the potential for clinical translation of these findings. Consistent with the RMC-4550 study, statistically significant antitumor efficacy was achieved in the U-CH1 mouse model treated with 20 mg/kg of TNO155 (*P* = 1.27×10^−6^), a clinical-stage SHP2 inhibitor with *ex vivo* anti-chordoma activity comparable to that of RMC-4550 and SHP099 (**Figure 5d, Supplementary Figures 6-7**).

TNO155 was also tested in two patient-derived xenograft (PDX) models, representing distinct patient populations of chordoma: CF539 was derived from a pediatric metastatic clival chordoma, and CF466 was derived from an adult metastatic lumbar chordoma. TNO155 treatment led to significant chordoma tumor growth suppression or regression in these models, with a good tolerability profile (**Figure 5d, Supplementary Figure 6**). Together, these findings using small-molecule perturbation of SHP2 further corroborate the essentiality of *PTPN11* in various models of chordoma, and thus nominate SHP2 inhibitors as a new therapeutic approach for the treatment of chordoma.

## DISCUSSION

Here we describe the emerging landscape of selective dependencies in chordoma. This study recovered the most significant known dependency gene in chordoma, *TBXT*, and further revealed a spectrum of novel selectively essential genes in this cancer type. These genes included several targetable or potentially targetable vulnerabilities that would not otherwise be detected by tumor sequencing approaches.

Collectively, the dependency genes identified herein can be classified into a broad range of biological functions. Canonical roles for these genes include: differentiation and development (the developmental transcription factors *TBXT, SOX9*, and *ZEB2*); proliferative signaling (*PTPN11* and *EGFR*); environmental sensing and metabolism (the aryl hydrocarbon receptor *AHR* and its transcription partner *ARNT*, the glucose transporter *SLC2A1*, the amino-acid transporter *SLC7A5*, the cAMP signaling regulatory component *PRKAR1A*, and the cholesterol metabolism regulator *UBIAD1*); cell-cycle progression (the cell cycle cyclin-dependent kinase *CDK6*, a cell-cycle gene regulator *THAP1*, and regulator of cell-cycle checkpoint activation and DNA replication *DSCC1*)^48,49^; immune regulation (*ADAR, PRKRA, PTPN11*, and a suppressor of the innate immune system *OTUD5*); RNA splicing (the U1 snRNP subunit gene *LUC7L2*, and a pre-mRNA splicing co-activator *SRRM2*)^50^; cellular transport (*SLC2A1, SLC7A5*, the ribosomal protein transport-related gene *HEATR3*, and a regulator of endoplasmic reticulum and secretory function *IER3IP1*)^51,52^; and DNA replication and repair (the Fanconi anemia core complex component *FANCM*, and *DSCC1*) (**Figure 6**).

**Figure 6.**
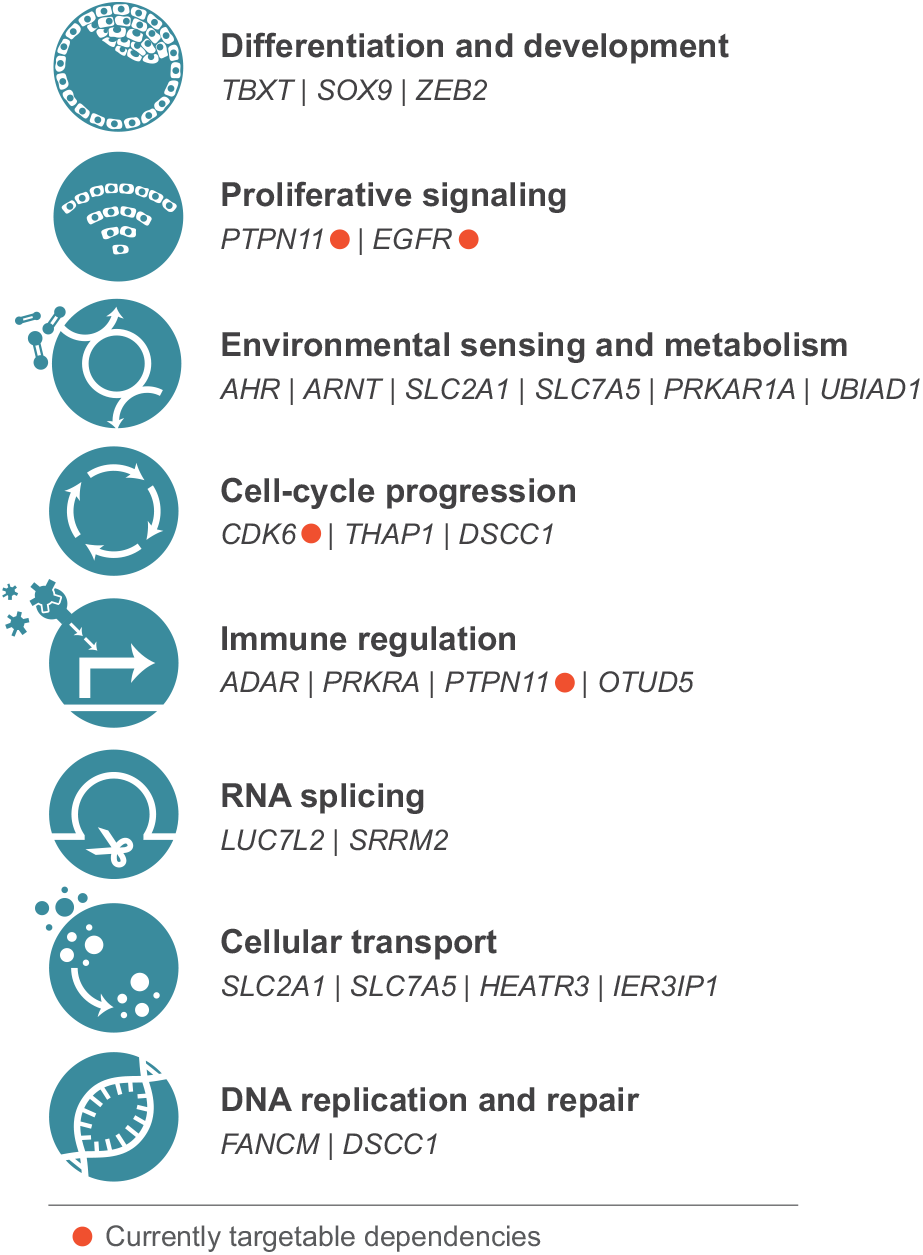
Functional classification of chordoma dependency genes.

We observed several instances of non-oncogene dependencies, whereby the unique cellular features of the cancer cell confer a heightened dependence on the normal function of specific genes^12^. For example, chordoma cells’ high intrinsic expression of ISGs appears to underlie their dependence on the double-strand RNA (dsRNA)-editing enzyme *ADAR*. It has been proposed that a chronic ISG state confers dependence on *ADAR* because it poises nucleic acid sensors like PKR to respond to the accumulation of endogenous immunogenic dsRNAs that *ADAR* normally edits^36,37,39^. Further investigation is needed to understand what triggers chronic ISG expression in chordoma. In other cancer types, the STING cytosolic DNA sensing pathway has been implicated in IFN-induced ISG expression, suggesting a link between chronic ISG expression and genomic instability^37,53^. Notably, recent evidence suggests that chordoma tumors frequently exhibit characteristics of defective homologous recombination DNA repair and increased genomic instability^54^; and some chordoma tumors harbor large numbers of clustered genomic rearrangements consistent with the phenomenon of chromothripsis^16,55^, features each of which is associated with the accumulation of immunostimulatory DNA^53,56^. It remains to be determined whether these elements contribute to chordoma cells’ high expression of ISGs and consequent dependence on *ADAR*.

Another dependency gene whose essentiality correlates with a specific cellular feature is *LUC7L2*. Here, dependency is correlated with lower copy number of a region containing the *LUC7L2* paralog *LUC7L*. This interaction is suggestive of a paralog lethality, whereby loss of one member of a paralog pair is associated with increased reliance on the other member^7,8^. These findings are consistent with that of a recent report demonstrating cross-regulation and partial redundancy between *LUC7L2* and *LUC7L*^57^.

Additionally, two genes scoring in our screens (*TBXT, SOX9*) can be classified as ‘lineage-survival’ dependency genes: these transcription factors mediate normal development of the embryonic notochord, the cell type from which chordoma is hypothesized to originate^58-60^. We also noted that three of the genes identified as chordoma dependency genes (*THAP1, SLC2A1, PRKRA*) are causative for dystonia, a neurological condition characterized by involuntary muscle contractions^61^. Further investigation is required to determine whether these results indicate a related etiology and/or common cell lineage from which chordoma and dystonia arise.

Lastly, our findings nominate an immediately actionable therapeutic target, SHP2, for the treatment of chordoma. Previous work has shown that cell lines dependent on *PTPN11* are most dependent on *EGFR*^46^. Our screens in chordoma similarly demonstrate dependency on both genes, but we noted that *PTPN11* dependency scores are of a greater magnitude than those of *EGFR* (**Figure 1, Supplementary Figure 1**). It is possible that SHP2 inhibition attenuates signaling mediated by different growth factor receptors, several of which can be activated concurrently in chordoma^62^. First-generation SHP2 inhibitors like TNO155 are currently under clinical evaluation (*e*.*g*., ClinicalTrials.gov identifier: NCT03114319); chordoma cells’ differential sensitivity to SHP2 suppression relative to other cancer types supports testing these inhibitors in chordoma patients.

## SUPPLEMENTARY FIGURE LEGENDS

**Supplementary Figure 1.**
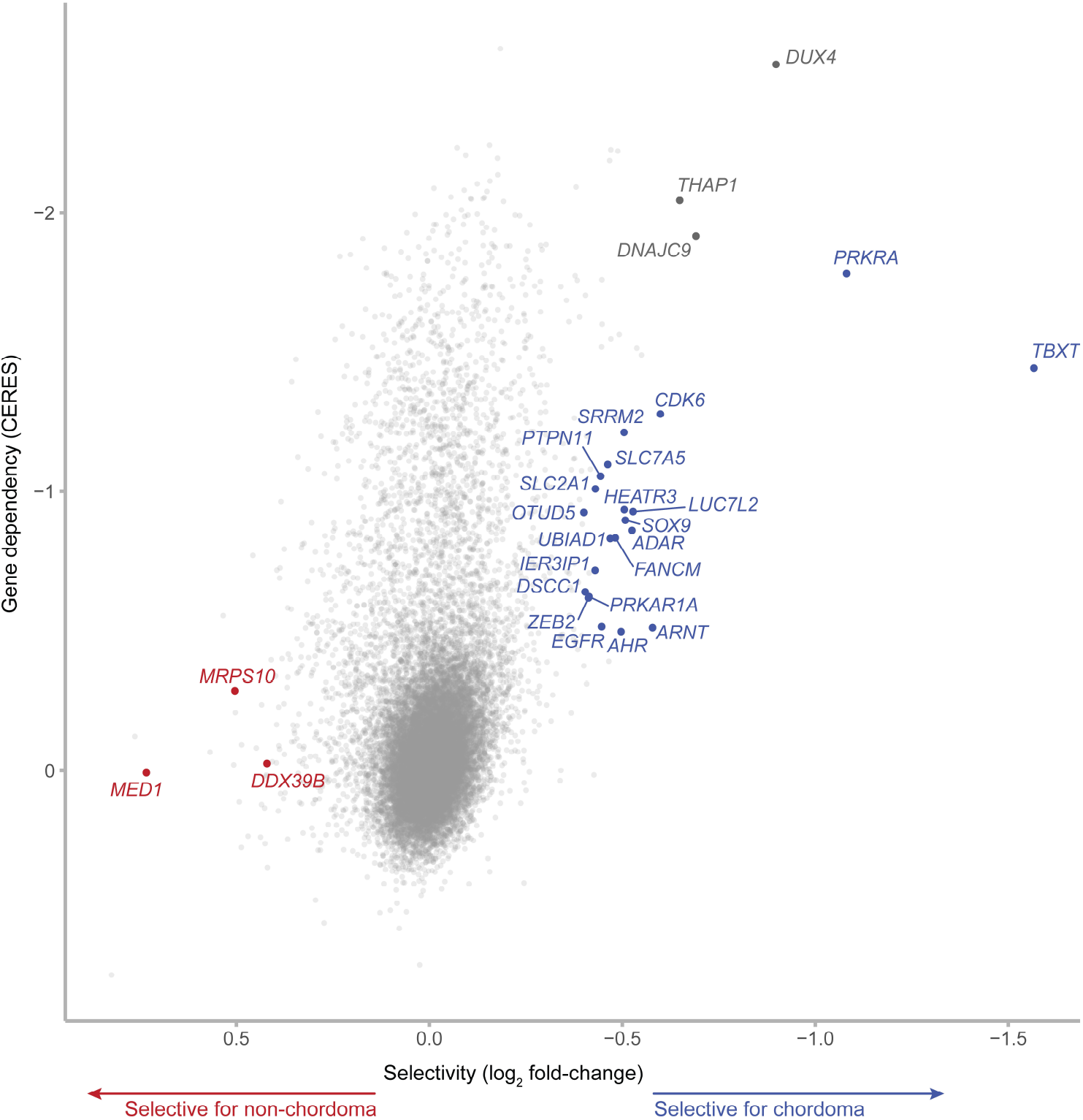
Genome-scale CRISPR-Cas9 screens identify a spectrum of selectively essential genes in chordoma (CERES analysis). Selective essentiality analysis identifying chordoma dependency genes. Selectivity is quantified by the log2 fold-change in mean gene dependency (using CERES scores) between four chordoma and 765 non-chordoma cell lines. Gene dependencies selective for chordoma/non-chordoma are indicated in blue/red (see Methods for details). Genes that do not meet the selectivity threshold for dependency probability scores (log2 fold-change > 0.3) but are strongly selective for chordoma based on CERES scores only (log2 fold-change > 0.5) are indicated and labeled in dark gray. They represent commonly essential genes that show a higher degree of viability effects in chordoma cell lines compared to non-chordoma cell lines.

**Supplementary Figure 2.**
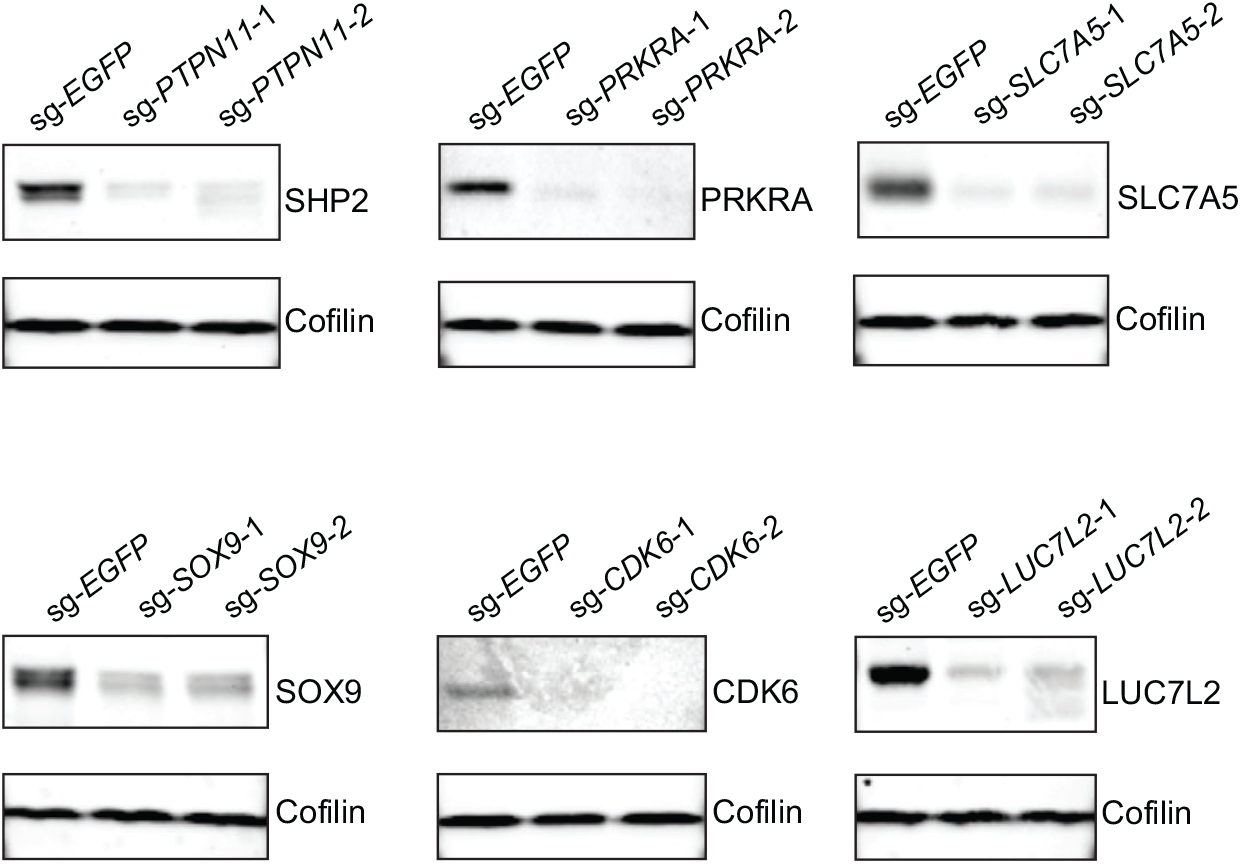
Immunoblot analysis confirming sgRNA-mediated protein repression. Immunoblot analysis of Cas9-expressing UM-Chor1 chordoma cells transduced with sgRNAs targeting a candidate dependency gene or a non-targeting sgRNA control.

**Supplementary Figure 3.**
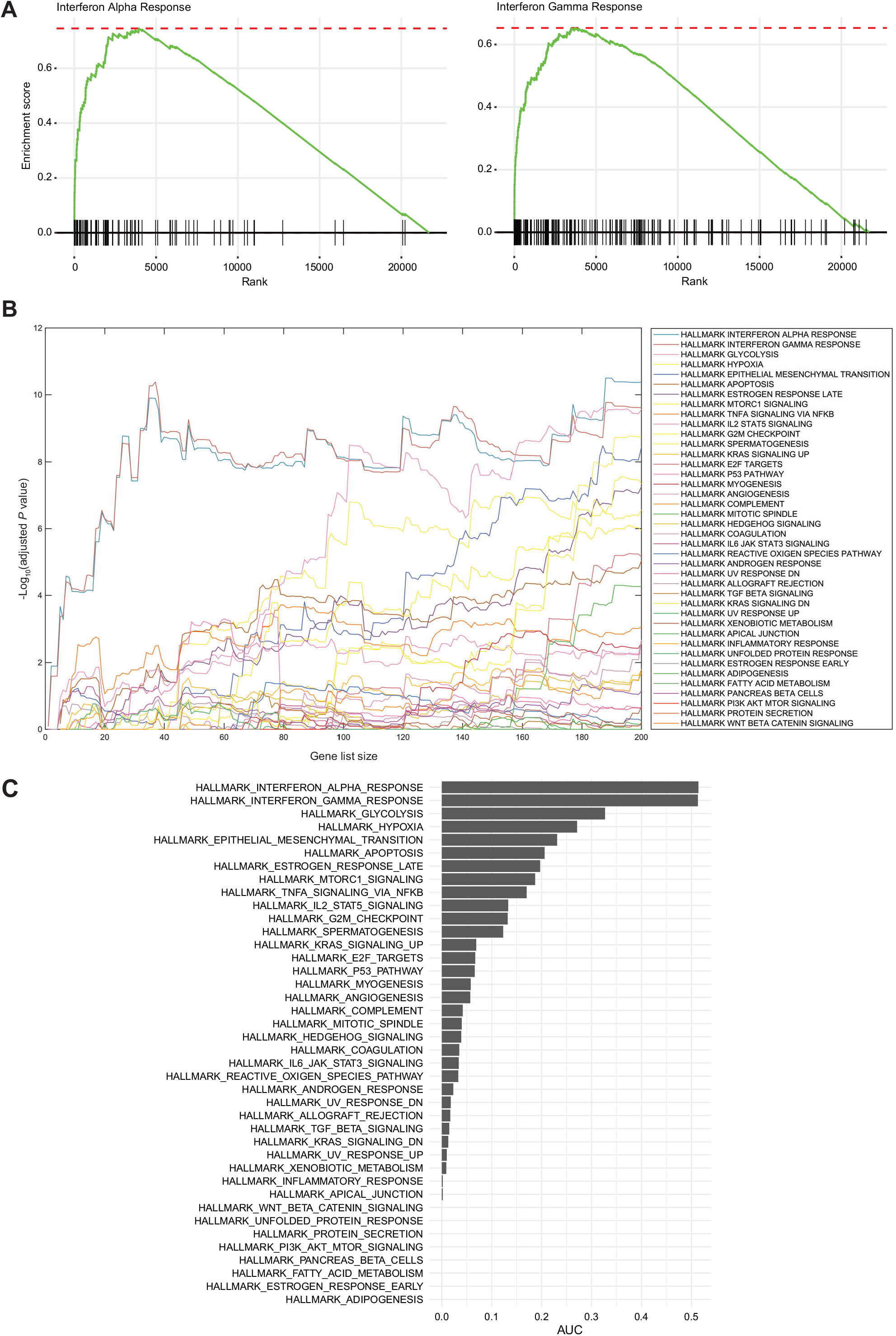
Interferon alpha and interferon gamma response genes are significantly upregulated following *ADAR* gene suppression in chordoma. **A)** Gene-set enrichment analysis (GSEA) plots for the two most-enriched gene sets from the hallmarks collection, Interferon Alpha (94 genes, enrichment score = 0.75, normalized enrichment score = 2.34, adjusted *P* = 3.84×10^−18^) and Interferon Gamma (188 genes, enrichment score = 0.65, normalized enrichment score = 2.16, adjusted *P* = 9.53×10^−20^). **B)** Adjusted GeLiNEA enrichment *P* values for all tested rank list sizes and gene sets. **C)** AUC values for GeLiNEA enrichment results.

**Supplementary Figure 4.**
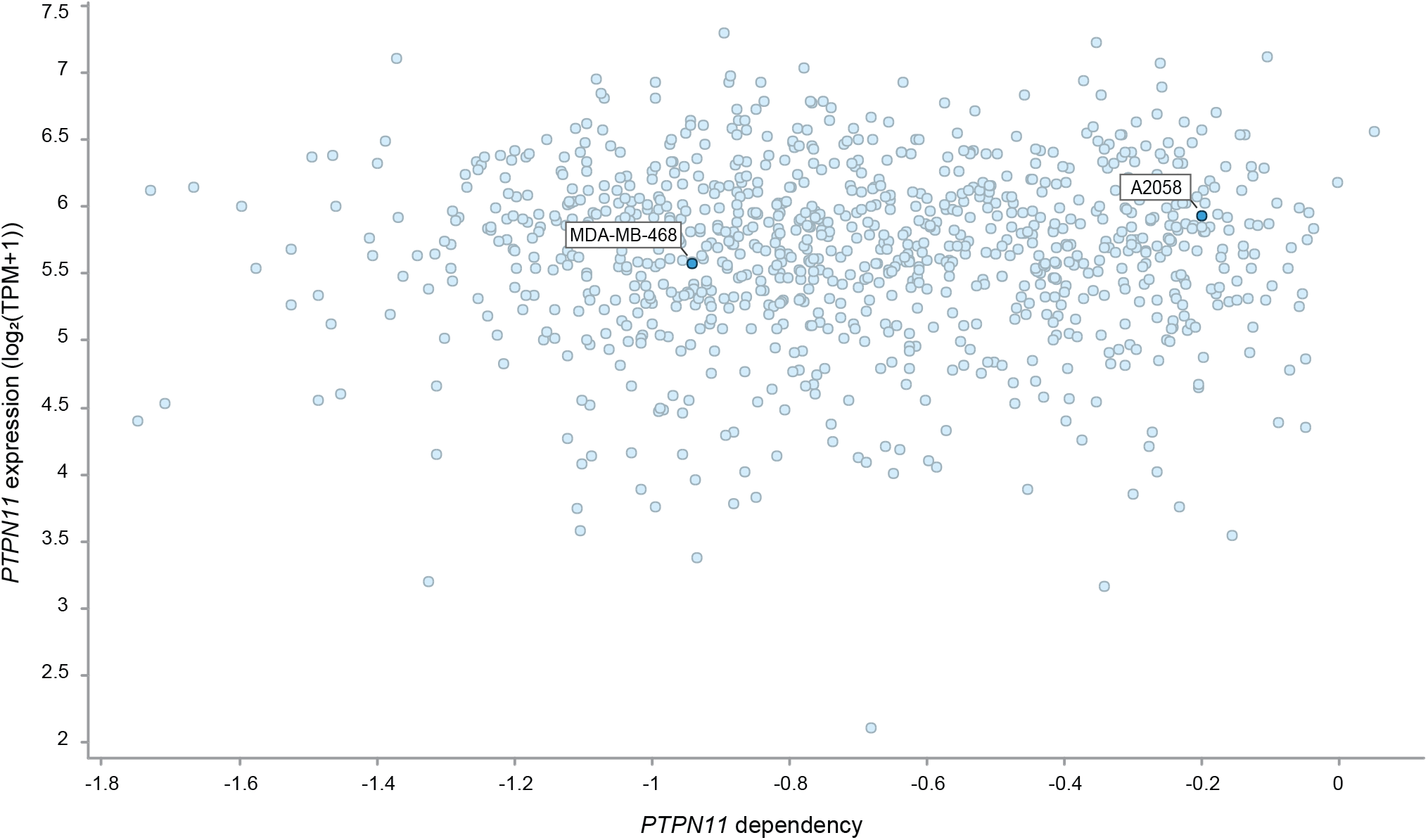
MDA-MB-468 and A2058 cell lines are sensitive and insensitive to loss of *PTPN11*, respectively. Data corresponding to *PTPN11* gene effect by CRISPR and *PTPN11* log2(TPM+1) expression for 952 cancer cell lines generated as part of the Broad Cancer Dependency Map. Gene effect is reported as a Chronos score^63^.

**Supplementary Figure 5.**
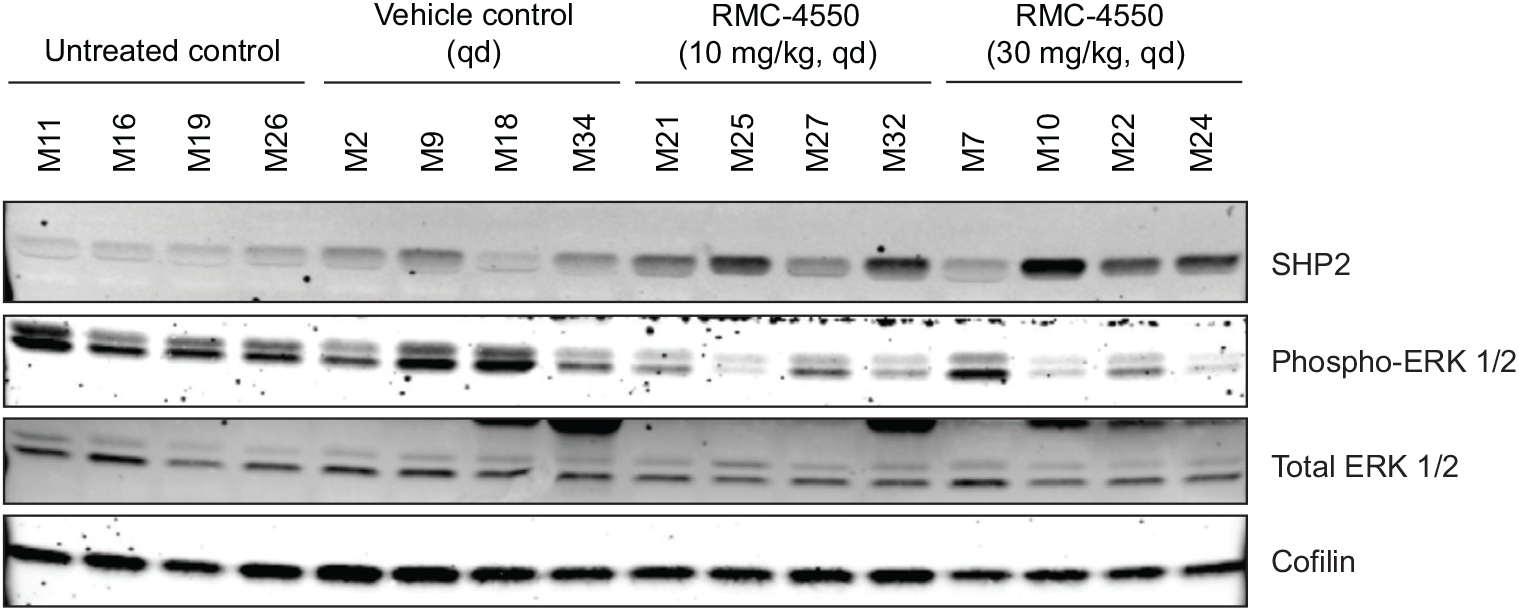
RMC-4550 treatment can reduce phosphorylation of ERK 1/2 *in vivo*. Immunoblot analysis of U-CH1 xenograft tumors following treatment with indicated doses of RMC-4550 once daily for 3 d.

**Supplementary Figure 6.**
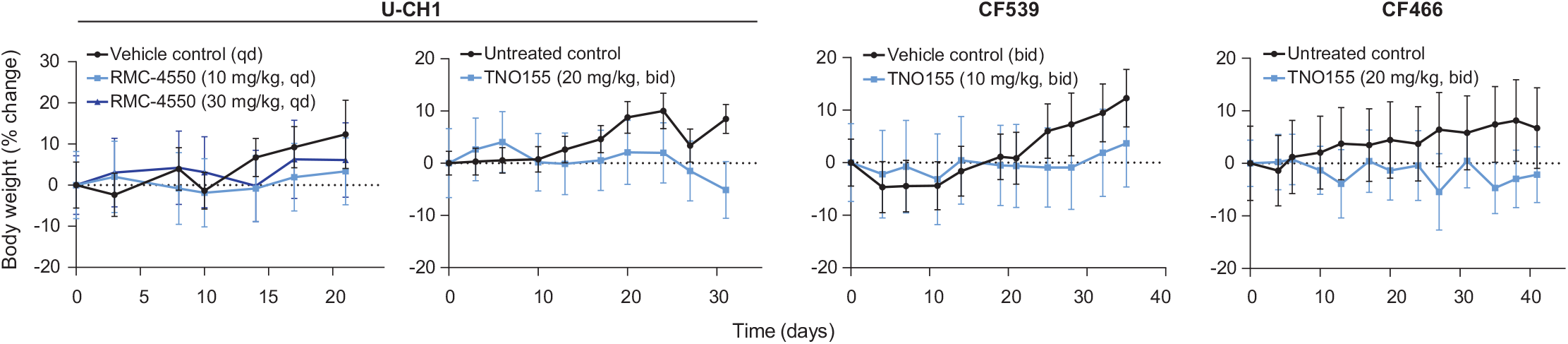
Tolerability of SHP2 inhibitor-treatment in mouse models of chordoma. Body weight (percent change relative to day 0 measurement) of mice engrafted with chordoma cells (U-CH1 cell line-derived xenograft, CF539 PDX, or CF466 PDX) and treated with a SHP2 inhibitor (RMC-4550 or TNO155). Points represent the mean body weight percent change ± s.e.m. (*n* = 4-5 tumors for each arm of the U-CH1/RMC-4550 study; 6-7 tumors for each arm of the U-CH1/TNO155 study; 6-7 tumors for each arm of the CF539 study; 7 tumors for each arm of the CF466 study).

**Supplementary Figure 7.**
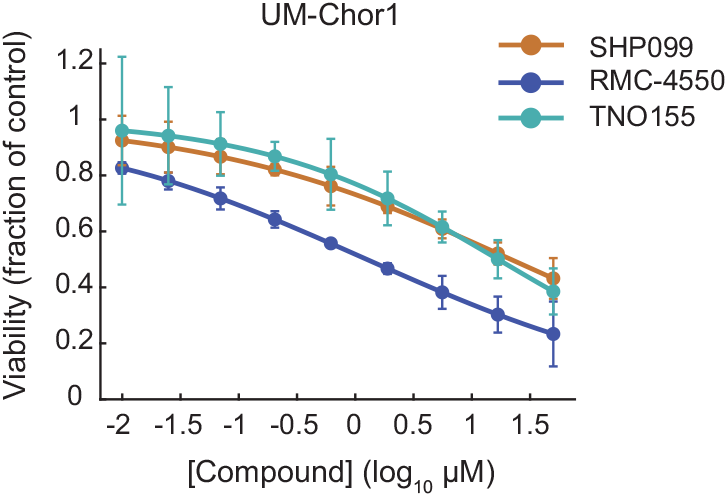
TNO155 has comparable potency to SHP099 and RMC-4550 in chordoma cells. Viability of UM-Chor1 cells treated with indicated concentrations of SHP2 inhibitors SHP099, RMC-4550, TNO155, or vehicle and assayed for cell viability after 6 d with CellTiter-Glo. Response data are represented by a fitted curve to the mean fractional viability at each concentration relative to vehicle-treated cells; error bars represent the s.e.m. (*n* = 4 biological samples measured in parallel).

## METHODS

### Cell lines and reagents

UM-Chor1, JHC7, MUG-Chor1, U-CH1, and U-CH2 chordoma cell lines were obtained from the Chordoma Foundation. CH22 cells have been described previously^64^ and were obtained from the Chordoma Foundation and Massachusetts General Hospital. MDA-MB-468 breast adenocarcinoma and A2058 melanoma cell lines were obtained from ATCC. UM-Chor1 cells were maintained in IMDM/RPMI (4:1) media + 10% fetal bovine serum (FBS) and 1X non-essential amino acids. JHC7 cells were maintained in DMEM/F12 (1:1) + 10% FBS. MUG-Chor1, U-CH1, and U-CH2 cell lines were maintained in IMDM/RPMI (4:1) media + 10% FBS. CH22 and MDA-MB-468 cells were maintained in RPMI media + 10% FBS. A2058 cells were maintained in DMEM media + 10% FBS. All chordoma cell lines were maintained on collagen I-coated plates.

RMC-4550 and SHP099 were purchased from MedChemExpress. TNO155 used for *ex vivo* studies was purchased from Selleck Chemicals and that used for *in vivo* studies was purchased from MedChemExpress.

### Genome-scale CRISPR-Cas9 screening

Genome-scale CRISPR-Cas9 screens using UM-Chor1 and MUG-Chor1 cell lines were performed previously^17^.

For U-CH2 and JHC7 cell lines, Cas9-expressing cells were generated as follows: each parental cell line was incubated with lentivirus corresponding to the pLX_311-Cas9 plasmid (Addgene plasmid #96924), encoding the Cas9 protein, in the presence of 4 μg/mL polybrene, dispensed in 12-well plates (1.5 × 10^6^ cells/well), and spin-infected at 2,000 rpm for 2 h at 30°C. After spin-infection, 2 mL of standard growth media was added to each well, and cells were incubated at 37°C overnight. The following day, for each cell line, cells were trypsinized and expanded in selective media containing 2-3 µg/mL blasticidin. Following selection for infected cells, Cas9 activity was confirmed in each transduced cell line using a Cas9-activity assay that has been described previously^65^.

Genome-scale screens were performed using a library of 74,687 unique sgRNAs targeting ∼18,560 genes (typically 4 sgRNAs per gene) and 1,000 non-targeting control sgRNAs (Broad Institute Avana sgRNA library)^18^. U-CH2-Cas9 and JHC7-Cas9 cells were each incubated with lentivirus corresponding to the pooled CRISPR library in the presence of 4 μg/mL polybrene, dispensed in 12-well plates (1.5 × 10^6^ cells/well across 21 plates for U-CH2 and 24 plates for JHC7), and spin-infected at 2,000 rpm for 2 h at 30°C. Lentivirus was titered in each cell line to achieve a low MOI (<1), and infections were performed with a sufficient a number of cells to achieve a representation of >700 cells per sgRNA per replicate after selection for infected cells. Following spin-infection, 2 mL of standard growth media was added to each well, and cells were incubated at 37°C overnight. The next day, for each cell line, cells were trypsinized, divided into three replicates, and expanded in selective media containing puromycin (14 µg/mL for U-CH2 and 5 µg/mL for JHC7) and blasticidin (3 µg/mL for U-CH2 and 1 µg/mL for JHC7). Cells were grown in culture for 20 d (U-CH2) or 22 d (JHC7) post-infection, with carryover of 5 × 10^7^ (U-CH2) or 4 × 10^7^ (JHC7) cells at each passage. Cells were grown in selective media until 8 d (JHC7) or 13 d (U-CH2) post-infection, after which they were grown in standard growth media. At 20 d (U-CH2) or 22 d (JHC7) post-infection, cells were collected and stored at −20°C in PBS until genomic DNA isolation steps.

Genomic DNA was isolated from cell pellets using the NucleoSpin Blood XL Kit (Macherey-Nagel). The sgRNA sequence was amplified by PCR with sufficient gDNA to maintain representation, and then quantified using massively parallel sequencing^66,67^. For each cell line, primary screening was performed once with three replicates.

### Computational analysis software for genetic studies

Unless otherwise stated, all genetic-perturbation analyses were performed in R (version 4.0.2) using the tidyverse package (version 1.3.0).

### Analysis of genome-scale CRISPR-Cas9 screening data

Chordoma screening data were pooled with other Project Achilles cell line data (https://depmap.org/portal/achilles/) and are available in public Broad Institute Cancer Dependency Map (DepMap) datasets (https://depmap.org/portal/). Additional details about the Project Achilles data processing pipeline can be found in a previous study^19^.

For all cell lines, CERES scores, dependency probability scores, and cell line annotations were obtained from the 20Q2 DepMap release (available at https://depmap.org/portal/download/). Further, for all non-chordoma cell lines, genomic and transcriptomic data (gene-expression values, copy-number variations, and mutations) were obtained from the same source. Genome and transcriptome sequencing data for chordoma cell lines were provided by the Chordoma Foundation (available at www.cavatica.org). Transcriptomic data for chordoma cell lines were processed as described below. Genomic data for chordoma cell lines were processed by and shared through the DepMap portal (see DepMap release notes for details; https://depmap.org). For this study, we used chordoma genomic data from the 20Q2 DepMap release, excepting mutation calls for U-CH2, which were obtained from the 22Q1 DepMap release.

To identify selective dependencies in chordoma, we compared gene-dependency scores between the four chordoma lines and 765 non-chordoma cell lines in the 20Q2 DepMap release using a linear model implemented in the R package limma (version 3.44.3)^68^. We performed this differential essentiality analysis for both CERES and dependency probability scores. The difference in mean dependency between the chordoma and non-chordoma lines was evaluated per gene as a log2 fold-change, and associated *P* values were derived from empirical-Bayes-moderated t-statistics. *Q* values were computed using the Benjamini– Hochberg method^69^.

Based on this analysis, selective dependencies were nominated using both CERES and probability score statistics. Genes were considered selective if (1) their CERES differential *P* value was lower than 0.02, (2) their absolute log2 fold-change in CERES scores exceeded 0.4, and (3) their absolute log2 fold-change in dependency probability exceeded 0.3. In addition, genes selective for chordoma needed to exceed an average dependency probability for chordoma cell lines of 0.5 and have a positive log2 fold-change (chordoma vs. non-chordoma). By contrast, genes selective for other cancer types needed to have an average dependency probability for chordoma cell lines equal to or lower than 0.5, with a negative log2 fold-change (chordoma vs. non-chordoma).

### Lentiviral vectors used for screening validation and functional characterization

To validate primary screening results, for each gene of interest, two sgRNA sequences represented in the Broad Institute Avana sgRNA library were selected and cloned into the lentiGuide-Puro plasmid (Addgene #52963). The spacer sequence for sg-*EGFP* has been described previously (“EGFP sgRNA 6”)^70^.

Spacer sequences for sgRNAs were as follows:

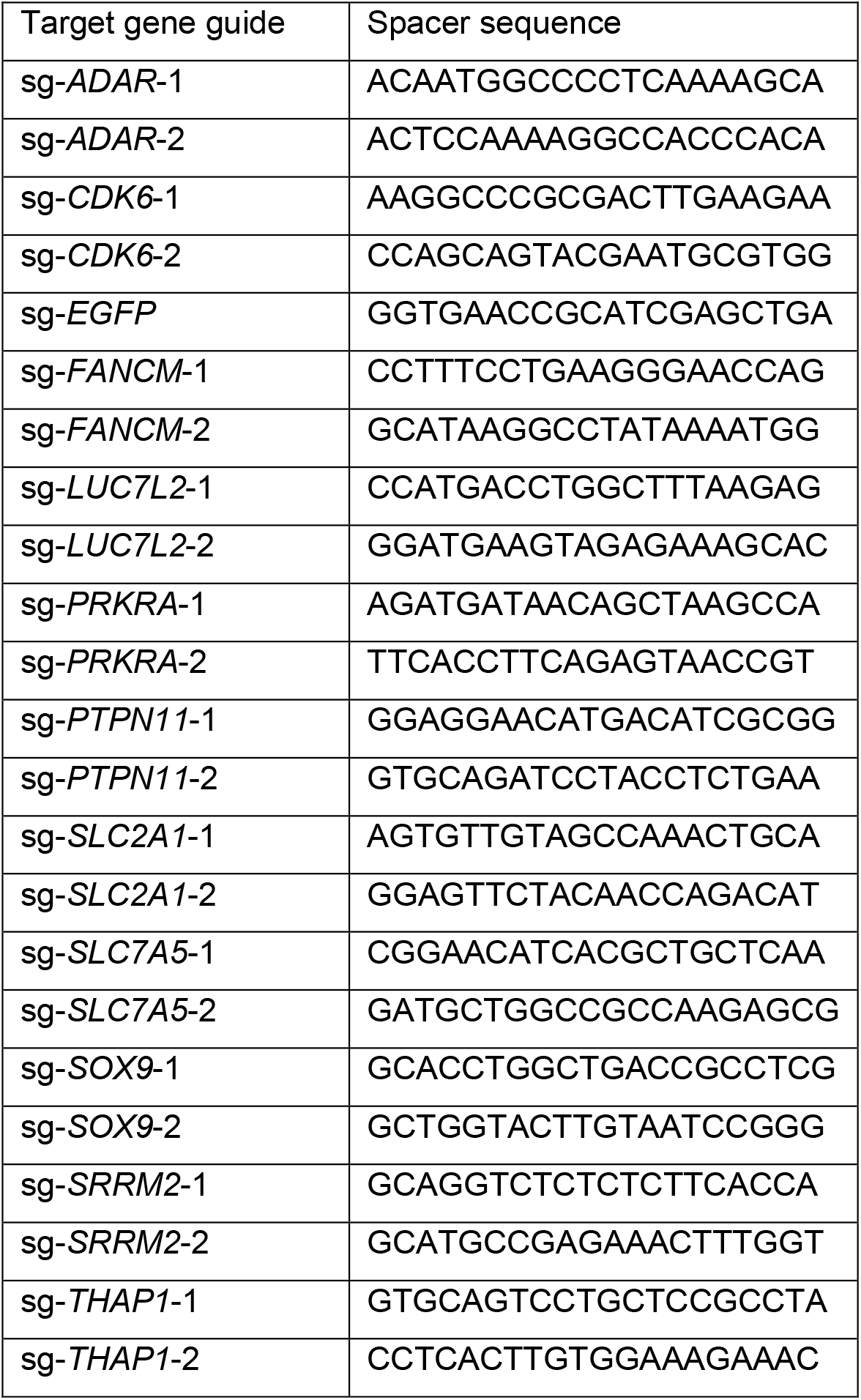

Lentivirus was produced by transfection of 293 T packaging cells with three plasmids (lentiGuide-Puro-sgRNA, psPAX2, and pMD2.G); and the Lipofectamine 2000 transfection reagent (Invitrogen). Media was changed to standard growth media the following day, and virus-containing supernatant was collected 3 d post-transfection.

### Validation of genome-scale CRISPR-Cas9 screens

UM-Chor1-Cas9 cells were generated as described previously^17^. UM-Chor1-Cas9 cells were seeded at a density of 300,000 cells/well in 6-well collagen I-coated plates. The next day, media was replaced with media containing lentivirus corresponding to the lentiGuide-Puro plasmid (Addgene #52963), encoding the sgRNA of interest (sequences below), in the presence of 8 µg/mL polybrene. Cells were spin-infected at 2,000 rpm for 30 min at 30°C. Following spin-infection, media was replaced with standard growth media and cells were incubated at 37°C overnight. The next day (at least 24 h post-infection), media was replaced with selective media containing 1 µg/mL puromycin. Following selection for infected cells, cells were seeded in 24-well plates in media containing 1 µg/mL puromycin at a density of 30,000 cells/well (three replicates per timepoint), and counted at indicated intervals over a 10 d period. Proliferation experiments were done four times for guides targeting *PTPN11, CDK6*, and *SLC7A5*; and twice for guides targeting all other genes, with minor variations in time intervals used for counting. Statistical analyses were performed with GraphPad Prism 9.

For immunoblotting to confirm sgRNA-mediated protein reduction, cells were transduced as above and selected cells were harvested 7 d post-transduction. Cell pellets were subjected to immunoblotting as described below. The experiment was performed three times for *PTPN11*; twice for *SLC7A5, PRKRA, LUC7L2*; and once for other genes.

For amplicon sequencing to confirm sgRNA-mediated genomic editing, cells were transduced as above and selected cells were harvested 6 d post-transduction. Cell pellets were processed as described below. The experiment was performed once.

### Immunoblotting

Cell pellets were resuspended in lysis buffer (50 mM Tris pH 7.4, 2.5 mM EDTA pH 8, 150 mM NaCl, 1% Triton X-100, 0.25% IGEPAL CA-630) supplemented with protease inhibitors (Roche) and Phosphatase Inhibitor Mixtures I and II (Calbiochem). Lysates were incubated on ice for >2 min, then centrifuged for 2 min at 15,700*g*. Protein in the supernatants was quantified using a BCA Protein Assay Kit (Pierce), normalized, reduced and denatured. Protein samples were resolved using Tris-Glycine gels (Novex), then resolved protein was transferred to iBlot Transfer Stack nitrocellulose membranes (Novex). Membranes were probed with primary antibodies at 4 °C overnight. The antibodies against SHP2 (clone D50F2, no. 3397, 1:1000), total Erk1/2 (clone 3A7; no. 9107, 1:500), phospho-Erk1/2 (Thr202/Tyr204) (no. 9101, 1:500), CDK6 (clone D4S8S, no. 13331, 1:500), PRKRA/PACT (clone D9N6J; no. 13490; 1:1,000), SLC7A5/LAT1 (no. 5347, 1:1,000), cofilin (clone D3F9, no. 5175; 1:10,000), and SOX9 (clone D8G8H, no. 82630, 1:1,000) were purchased from Cell Signaling Technology. The antibody against LUC7L2 (no. PA5-62446, 1:1,000) was purchased from Invitrogen. Membranes were incubated with IRDye secondary antibodies (1:10,000; LI-COR Biosciences) and detected with the Odyssey Imaging System (LI-COR Biosciences).

### Amplicon sequencing of sgRNA-mediated genomic edit sites

Genomic DNA was purified from cell pellets using the QIAamp DNA Mini Kit (Qiagen). A 200-275 base pair region containing the relevant sgRNA target sequence was PCR-amplified from genomic DNA using the NEBNext High-Fidelity 2X PCR Master Mix (New England BioLabs) in a 25 µL PCR reaction volume (primer sequences appear below). PCR products were purified using the QIAquick PCR purification Kit (Qiagen).

Library preparation and sequencing were performed at the Dana-Farber Cancer Institute Molecular Biology Core Facilities. cDNA amplicons were fragmented to approximately 250 base pairs using Covaris adaptive focused acoustics on the M220 platform. Illumina sequencing libraries were prepared using Swift S2 Acel reagents on a Biomek i7 liquid handling platform. Finished libraries were quantified by Qubit fluorometer, Agilent TapeStation 2200, and RT-qPCR using the Kapa Biosystems library quantification kit according to manufacturer’s protocols. Uniquely indexed libraries were pooled in equimolar ratios and sequenced on an Illumina MiSeq with paired-end 150-base pair reads.

Amplicon sequencing was performed once.

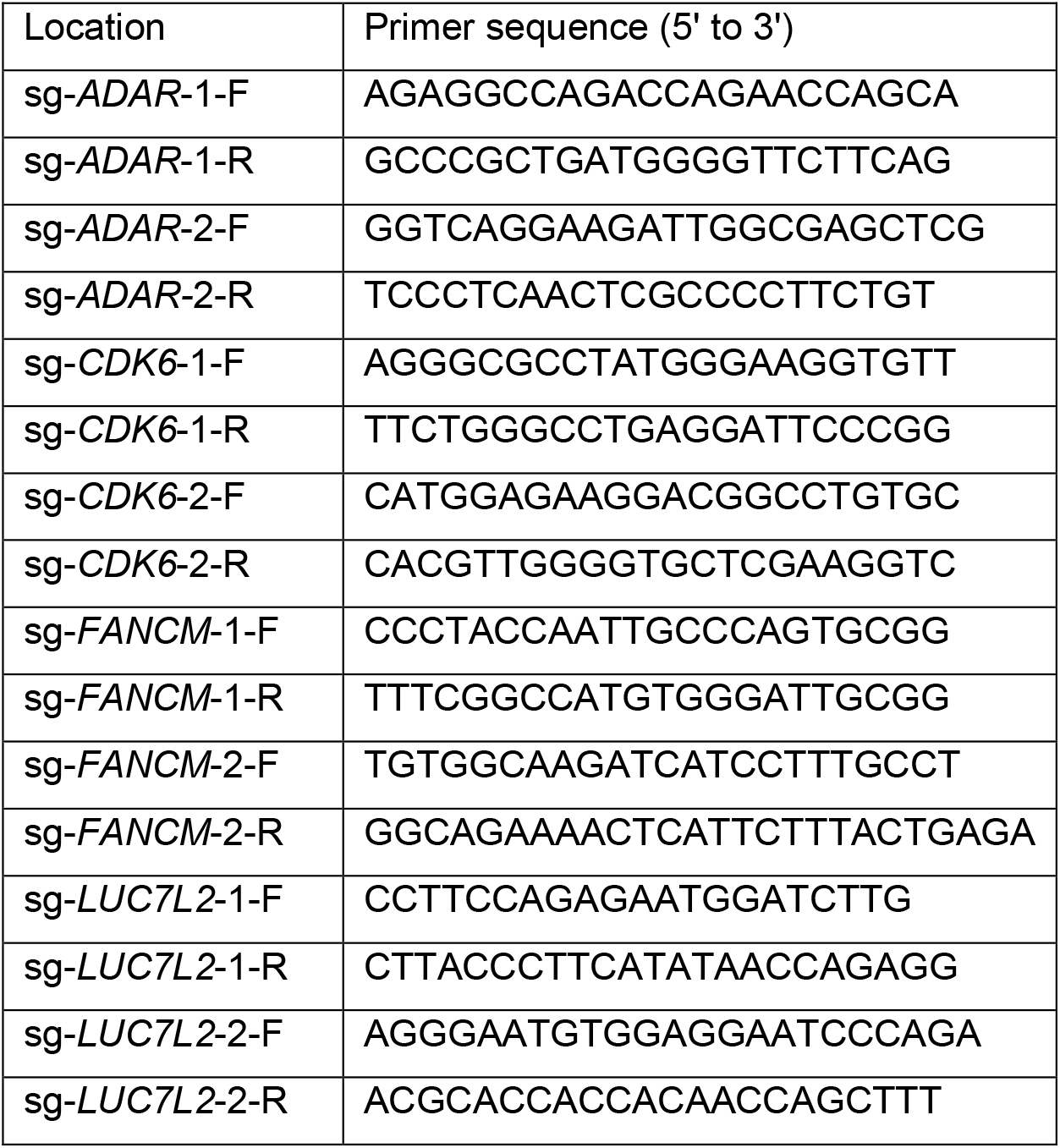
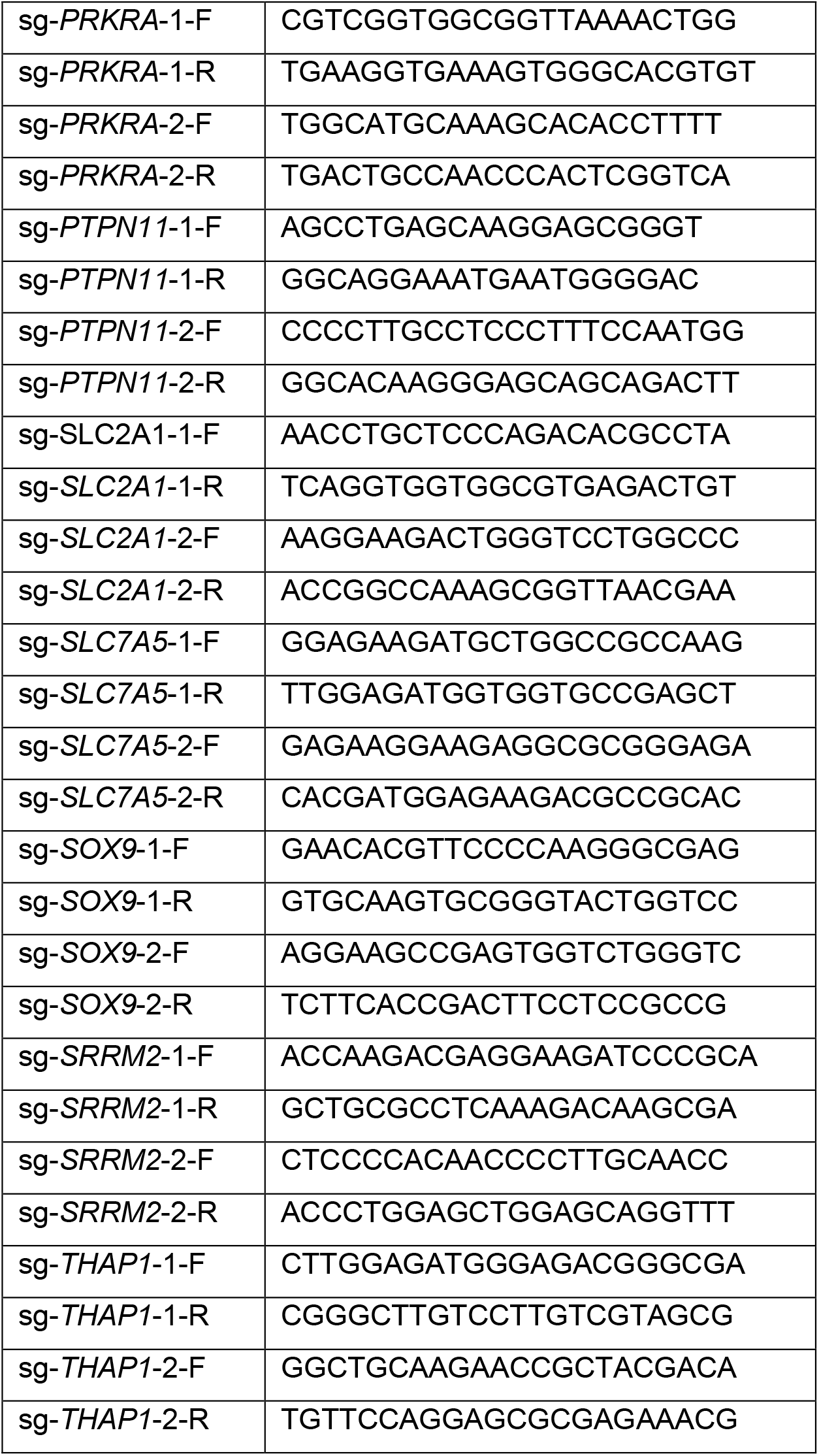

### Amplicon sequencing analysis

We evaluated the editing performance of sgRNAs from amplicon sequencing data using CRISPResso2 (version 2.1.3)^71^ with default parameters. Gene reference sequences were obtained from the NCBI Nucleotide database using the R package rentrez (version 1.2.3) and expected amplicon sequences extracted with the matchProbePair function from the R package Biostrings (version 2.56.0). Potential off-target genomic sequences were identified with NCBI BLAST (https://blast.ncbi.nlm.nih.gov/Blast.cgi) and provided to CRISPResso2 as alternate sequences to reduce artifactual reporting of editing.

### Pathway analysis of chordoma dependency genes

#### Co-essentiality network

The network was generated using the R package igraph (version 1.2.6). First, all selective chordoma dependency genes were added as nodes. To draw edges between gene pairs, we calculated the Pearson correlation coefficient between their dependency probability score profiles (dependency probability scores for all 769 cell lines). Gene pairs with a correlation of least 0.18 were connected by an edge. We visualized the resulting network in Cytoscape (version 3.8.2), coloring nodes by gene-dependency probability scores and scaling edge width by Pearson correlation coefficients.

#### Protein-protein interaction network

We performed a multi-protein search in STRING (version 11.5) (https://string-db.org/)^72^ with default parameters, using HGNC symbols for all selective chordoma dependency genes as input. The resulting network was exported and visualized in Cytoscape (version 3.8.2), coloring nodes by gene-dependency probability scores and scaling edge width by STRING confidence scores. Singletons, removed automatically during export from STRING, were manually re-added in Cytoscape.

### RNA-sequencing analysis of parental chordoma cells

Gene-expression levels of chordoma cell lines were quantified from RNA-seq data using the DepMap RNA-seq pipeline (https://github.com/broadinstitute/depmap_omics) on the Terra computing platform (https://terra.bio/). Briefly, RNA-seq FASTQ files were aligned to the hg38 reference genome with STAR (v2.5.3a)^73^. Gene- and transcript-level expression was then quantified with RSEM (v1.3.0)^74^ to obtain transcript per million (TPM) values.

### Genomic and transcriptomic correlates of chordoma dependency genes

For all chordoma dependency genes, we calculated Pearson correlation coefficients between their CERES dependency profiles (CERES scores for a given gene across all cell lines) and genome-wide (1) gene-expression profiles (gene-level log2(TPM+1) values across all cell lines), (2) copy-number profiles (gene copy number across all cell lines), and (3) mutation profiles (gene mutations across all cell lines, indicating the presence or absence of mutations as 1 and 0, respectively). Correlates for each dependency were then ranked by decreasing correlation coefficients.

For dependencies correlated with gene-copy-number changes, we nominated potentially causal genes based on known biological connections to the dependency gene. We further tested whether the dependency also correlated with the copy number of genes surrounding the nominated gene on the same chromosome. To do so, we first selected all genes within windows ranging from 5×10^5^ bp to 5×10^6^ bp (step size: 5×10^5^) around the nominated gene. The optimal window was then determined using gene-set enrichment analysis^40^ via the fgsea function from the synonymous R package^75^ (settings: eps=0, default for all others). As input, we used (1) the gene sets obtained from each window size and (2) the list of all gene-copy-number correlates, ranked by their correlation with the dependency gene.

### ISG expression in chordoma and non-chordoma cancer cell lines

A previously described procedure^37^ was followed, using the ISG core signature from the same study (*ADAR, BST2, CASP1, CMPK2, CXCL10, DDX60, DHX58, EIF2AK2, EPSTI1, GBP4, HERC6, IFI35, IFIH1, IFIT2, IFIT3, IRF7, ISG15, ISG20, MX1, NMI, OASL, OGFR, PARP12, PARP14, PNPT1, PSME2, RSAD2, RTP4, SAMD9L, SP110, STAT2, TDRD7, TRAFD1, TRIM14, TRIM21, TRIM25, UBE2L6, USP18*). Briefly, mean absolute deviation modified z-scores (ZMAD) were calculated for each gene, using median and mean absolute deviation for TPM values across all cell lines. The ISG core score was then calculated as the mean ZMAD across all ISG signature genes. Cell lines without disease annotation, disease annotations with only one cell line (teratoma and adrenal cancer), and the disease annotation “Engineered” were omitted from the analysis.

### RNA sequencing and analysis of sgRNA-treated cells

UM-Chor1-Cas9 cells were seeded at a density of 300,000 cells/well in 6-well collagen I-coated plates. Cells were transduced with lentivirus and selected for infected cells as described in the Methods section corresponding to CRISPR-Cas9 screening validation. Following selection for infected cells, selective growth media was replaced with standard growth media. One day post-media change (6 d post-transduction), cells were harvested. Total RNA was extracted from cells using an RNeasy kit (Qiagen).

Library preparation, sequencing, and sequencing analysis were performed at the Dana-Farber Cancer Institute Molecular Biology Core Facilities. Libraries were prepared using Roche Kapa mRNA HyperPrep strand specific sample preparation kits from 200 ng of purified total RNA according to the manufacturer’s protocol on a Beckman Coulter Biomek i7. The finished dsDNA libraries were quantified by Qubit fluorometer and Agilent TapeStation 4200. Uniquely dual indexed libraries were pooled in an equimolar ratio and shallowly sequenced on an Illumina MiSeq to further evaluate library quality and pool balance. The final pool was sequenced on an Illumina NovaSeq 6000 targeting 40 million 100-base pair read pairs per library.

Sequenced reads were aligned to the UCSC hg38 reference genome assembly, and gene counts were quantified using STAR (v2.7.3a)^73^. RNA-seq analysis was performed using the VIPER Snakemake pipeline^76^. Differential gene-expression testing was performed by DESeq2 (v1.22.1)^77^.

We performed gene-set enrichment analysis^40^ using the fgsea function from the synonymous R package^75^ (settings: eps=0, default for all others) on the MSigDB hallmark collection of gene sets (version 7.4)^78^, with log2 fold-changes from DESeq2 as input.

We used GeLiNEA to quantify enrichment while taking biological network information into account^41^. We ran GeLiNEA via the Molecular Data Provider (MolePro) API (https://translator.broadinstitute.org/molecular_data_provider/assets/lib/swagger-ui/index.html?url=/molecular_data_provider/assets/openapi.json#/), using the MSigDB hallmark collection of gene sets and the STRING interaction network (version 10.5, *Homo sapiens*, including all available evidence types)^72^. The source code for GeLiNEA can be obtained at https://github.com/broadinstitute/GeLiNEA. The API calls were made using a MATLAB script (2018b version), with the parallelization toolbox for parallelizing the calls.

Rather than a ranked list, GeLiNEA requires a list of top-scoring genes as input (in addition to a list of known gene sets). To account for differences in list size, we considered all lists from 1 to 200 top-scoring genes, with gene ranks determined by their adjusted *P* value from the previously described DESeq2 differential-expression analysis. Using a MATLAB 2018b implementation, we then constructed curves charting gene list size against the negative log10 of adjusted enrichment *P* values, and ranked hallmark gene sets based on their normalized AUC, calculated as follows:

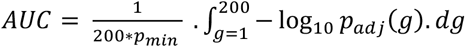

where 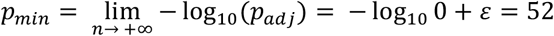, and ε equals the inverse logarithm base 10 of the floating-point relative accuracy.

RNA sequencing was performed once.

### Interferon-β ELISA

UM-Chor1-Cas9 cells were seeded at a density of 300,000 cells/well in 6-well collagen I-coated plates. Cells were transduced with lentivirus and selected for infected cells as described in the Methods section corresponding to CRISPR-Cas9 screening validation. Following selection for infected cells, selective growth media was replaced with standard growth media. At one and two days post-media change, an aliquot of conditioned media was harvested from cells and centrifuged to remove any residual cells. The supernatant was assayed for interferon-β levels using the VeriKine-HS™ Human IFN Beta Serum ELISA Kit (PBL Assay Science), following ‘Protocol A’ provided by the manufacturer.

Absorbance values of the ELISA were measured at 450 nm using a SpectraMax M5 microplate reader (Molecular Devices). To calculate interferon-β titers, optical densities (ODs) for media alone samples were first subtracted from the standard and sample ODs to eliminate background, and the interferon-β concentration in the samples was determined from a standard curve, fit with a sigmoidal, 4-parameter logistic equation (GraphPad Prism 9). Statistical analyses were performed with GraphPad Prism 9. The conditioned media was collected once and the ELISA was performed three times on day 2 samples.

### Computational analysis software for small-molecule sensitivity studies

Computational analyses and visualizations for small-molecule studies were performed in Microsoft Excel, Pipeline Pilot (v 18.1.0.1604; Biovia, Corp.), or MATLAB 2018b (MathWorks, Inc.).

### High-throughput small-molecule sensitivity studies

U-CH1, U-CH2, JHC7, MUG-Chor1, UM-Chor1, CH22, MDA-MB-468, and A2058 cells were each seeded overnight in 384-well BioCoat Collagen I (Corning) microtiter plates at a density of 2,000, 1,600, 1,600, 1,800, 1,200, 1,200, 1,200, and 1000 cells per well, respectively. The following day, compound or DMSO was added to wells using an HP D300 digital dispenser instrument. Each compound was tested using nine concentrations, in quadruplicate (four wells treated in parallel). Cell viability was assayed 6 d after compound addition with the CellTiter-Glo reagent (Promega). Luminescence well values were normalized to DMSO-treated wells by subtracting per-plate average DMSO log2-luminescence values from the log2-luminescence values of each treatment well.

Data pre-processing from instrument files through DMSO normalization was performed in Pipeline Pilot except for the experiment depicted in Supplementary Figure 7, which was normalized in Microsoft Excel and MATLAB; curve-fitting, numerical integration, and subsequent analysis steps were performed in MATLAB. For all drug-printer experiments, curves were fit using all data points as inputs to curve-fitting and numerical integration. Curve fitting (to derive EC50 and other curve parameters) and numerical integration (to derive AUC) were otherwise performed as described previously^17^.

SHP099 was tested four times in UM-Chor1 cells (including in the experiment depicted in Supplementary Figure 7), twice in JHC7, and once in all other cell lines. RMC-4550 was tested three times in UM-Chor1 (including in the experiment depicted in Supplementary Figure 7), twice in JHC7, and once in all other cell lines. TNO155 was tested once in UM-Chor1 cells.

### Colony formation assays

U-CH1, U-CH2, JHC7, MUG-Chor1, UM-Chor1, CH22, MDA-MB-468, and A2058 cells were seeded in 6-well plates at a density of 70,000, 50,000, 80,000, 90,000, 18,000, 6,000, 15,000, and 2,000 cells/well, respectively. The following day, RMC-4550 or DMSO was added to wells at a 1:1000 dilution. Cells were cultured in compound- or DMSO-containing media for a total of 14 d, with compound- or DMSO-containing media replenished at 7 d post-treatment. At the experimental endpoint, compound- or DMSO-containing media was aspirated, and cells were first washed with PBS, then fixed with 100% methanol for 10 minutes. Methanol was aspirated and cells were stained with 0.5% crystal violet (Alfa Aesar) staining solution in 25% methanol for 10 minutes. Staining solution was aspirated and plates were washed with water and subsequently air dried. Each cell line was tested at least four times, with minor variations in cell seeding densities.

Plates were imaged with an Epson Perfection V550 Photo scanner.

### Immunoblots of compound-treated cell lines

For immunoblots of SHP2-inhibitor-treated cell lines: for each cell line, cells were seeded in a 6-well plate at a density of 400,000 cells/well. The following day, cells were treated with RMC-4550, SHP099, or DMSO (1:1000 dilution in media) for 2 h before being harvested. Cell pellets were lysed and processed for immunoblotting as described above. The experiment was performed at least twice for UM-Chor1 and JHC7, and once for all other cell lines.

### *PTPN11* dependency in MDA-MB-468 and A2058 cell lines

Data corresponding to *PTPN11* log2(TPM+1) expression (21Q3 Public) and *PTPN11* gene effect by CRISPR (DepMap 21Q3 Public+Score, Chronos) for 952 cancer cell lines were obtained from the Broad Institute Cancer Dependency Map Portal (https://depmap.org/portal/).

### Conditioned media assays

UM-Chor1-Cas9 cells were seeded at a density of 300,000 cells/well in 6-well collagen I-coated plates. Cells were transduced with lentivirus and selected for infected cells as described in the Methods section corresponding to CRISPR-Cas9 screening validation. Following selection for infected cells, selective growth media was replaced with standard growth media. Three days post-media change, conditioned media was harvested from cells and centrifuged to remove any residual cells. The supernatant was used to replace the standard growth media of UM-Chor1 cells previously seeded in 24-well plates (seeding density of 30,000 cells/well), with three replicate wells treated per supernatant condition. Cells were counted after three days of treatment with conditioned media. Statistical analyses were performed with GraphPad Prism 9. The experiment was performed twice.

### *In vivo* xenograft studies

Animal experiments were performed at XenoSTART in San Antonio, TX in tumor-bearing mice. Six- to twelve-week-old female athymic nude mice were implanted subcutaneously with low-passage tumor fragments. When tumors reached approximately 150-300 mm^3^ (for efficacy studies) or 300-500 mm^3^ (for the pharmacodynamics study), animals matched by tumor volume (TV) were randomized into control and treatment groups, each group containing 4-7 animals. RMC-4550 was formulated in 2% hydroxypropyl methylcellulose E-50, 0.5% Tween-80 in 50 mM sodium citrate buffer, pH 4.0. TNO155 was formulated in 0.5% Tween-80, 0.5% methylcellulose. Dosing began at Day 0 and drugs (or respective vehicles) were administered orally at the dose levels and schedules noted in the main text. Animals were observed daily, and weights and TVs were measured twice a week using an electronic scale and digital calipers, respectively. Tumor dimensions were converted to TV using the formula: TV (mm^3^) = width^2^ (mm) x length (mm) x 0.52. The study endpoints were when the control group reached mean TV = 1500 mm^3^ or a specified time point. For mice that reached tumor volume endpoint (one vehicle-treated mouse on day 25 of the CF539/TNO155 study), the final TV measurement was carried over and plotted for the remainder of the study, and body weight measurements were not plotted for this mouse beyond this date. Statistical analyses were performed with GraphPad Prism 9.

Immunoblotting of tumor tissue was performed by first homogenizing tumor fragments in lysis buffer (recipe described in the Immunoblotting methods section) using the Precellys Evolution instrument (Bertin Technologies) and the following protocol: 3 × 7,500 r.p.m. for 20 s, pausing for 15 s between rounds. Homogenized lysates were subsequently centrifuged for 2 min at 15,700g. Supernatants were quantified and subjected to immunoblotting as described in the Immunoblotting methods section. Immunoblots were performed twice.

## AUTHOR CONTRIBUTIONS

T.S. and S.L.S. designed and supervised the study. M.J.W. designed the computational analyses of genetic data. T.S., A.G., and Y.L. performed experiments. T.S. performed low-throughput genetic-perturbation experiments, small-molecule sensitivity experiments, RNA-sequencing experiments, and immunoblotting. A.G. and Y.L. performed genome-scale CRISPR-Cas9 screening with assistance from T.S. M.J.W., M.K., J.D., S.M., and P.A.C. performed computational analyses. M.J.W. performed co-essentiality/interaction network analysis, genomic/transcriptomic analysis, ISG-core-score analysis, and analysis of CRISPR amplicon sequencing data. M.J.W., M.K., and J.D. performed analysis of genome-scale CRISPR-Cas9 screening data. M.J.W., S.M., and J.K. performed analysis of RNA-sequencing data. P.A.C. performed analysis of small-molecule sensitivity profiling data. F.V., D.E.R., and W.C.H supervised genome-scale CRISPR-Cas9 screening studies. D.M.F., J.L., and J.S. supervised *in vivo* experiments, which were performed at XenoSTART. T.S. wrote the manuscript, which was revised by M.J.W. and S.L.S. All authors reviewed and/or provided feedback on the manuscript.

## ACKNOWLEDGEMENTS

The authors thank Mary O’Reilly and Elena Berg for assistance with creating illustrations; members of the Broad Institute Genetic Perturbation Platform for sgRNA library construction and technical assistance with DNA processing after genome-scale CRISPR-Cas9 screening; XenoSTART for conducting *in vivo* studies; the Dana-Farber Cancer Institute Molecular Biology Core Facilities for assistance with RNA and CRISPR amplicon sequencing; A. Cherniack for guidance with copy-number analysis; and J. Gasser for helpful discussions. This work was generously supported by the Chordoma Foundation, and the NCI’s Cancer Target Discovery and Development (CTD^2^) Network (grant number U01CA217848 awarded to S.L.S. and U01CA176058 awarded to W.C.H.).

## COMPETING INTERESTS

M.J.W. is an employee and shareholder of Kojin Therapeutics. F.V. receives research support from Novo Ventures. D.E.R. receives research funding from members of the Functional Genomics Consortium (Abbvie, Bristol-Myers Squibb, Jannsen, Merck, Vir), and is a director of Addgene, Inc. W.C.H. is a consultant for ThermoFisher, Solasta, MPM Capital, Frontier Medicines, Tyra Biosciences, RAPPTA Therapeutics, KSQ Therapeutics, Jubilant Therapeutics, Function Oncology and Calyx. P.A.C. is an advisor to Pfizer, Inc., and Belharra Therapeutics. S.L.S. serves on the Board of Directors of the Genomics Institute of the Novartis Research Foundation (‘GNF’); is a shareholder and serves on the Board of Directors of Jnana Therapeutics; is a shareholder of Forma Therapeutics; is a shareholder and advises Kojin Therapeutics, Kisbee Therapeutics, Decibel Therapeutics and Eikonizo Therapeutics; serves on the Scientific Advisory Boards of Eisai Co., Ltd., Ono Pharma Foundation, Exo Therapeutics and F-Prime Capital Partners; and is a Novartis Faculty Scholar. All other authors declare no competing interests.

## Notes

### Summary of Updates

Figures repositioned to improve readability.

